# Computational Modeling of Functionalized Graphene Quantum Dots Binding to Tau Fibrils in Alzheimer’s Disease

**DOI:** 10.64898/2026.07.29.741532

**Authors:** Max Walton-Raaby, Subha Kalyaanamoorthy

**Affiliations:** Department of Chemistry, University of Waterloo, Waterloo, Ontario, N2L 3G1, Canada; Waterloo Artificial Intelligence Institute, University of Waterloo, Waterloo, Ontario, N2L 3G1, Canada; Waterloo Institute of Nanotechnology, University of Waterloo, Waterloo, Ontario, N2L 3G1, Canada

**Keywords:** **Keywords:** Alzheimer’s disease, Tau protein, Tau aggregates, graphene quantum dots, nanomedicine, functionalization, molecular dynamics, protein-ligand interactions

## Abstract

The aggregation of Tau protein into straight filaments (SFs) and paired helical filaments (PHFs) is central to Alzheimer’s disease (AD) pathology and a key target for therapeutic inhibition. Graphene quantum dots (GQDs) are biocompatible nanomaterials that have shown promise in inhibiting amyloidogenic protein aggregation across related neurological pathologies. The effect of GQD functionalization on interactions with Tau aggregates (TAs) is poorly understood, though recent evidence suggests that anionic GQDs are effective TA inhibitors. In this study, we survey how GQD functionalization influences binding to SFs and PHFs to guide future development of therapeutic GQDs. We identify binding sites in SFs and PHFs, dock our GQD library to these sites, and perform molecular dynamics simulations on promising complexes, totaling 28 *µ*s of sampling. We discover that anionic GQDs preferentially bind to the positively charged SF large protofilament interface, whereas in PHFs, anionic GQDs have a modest binding preference for the C-shaped curve region. Binding of GQDs at the C-shaped curve in both TAs induces distinct protofilament conformational dynamics resembling a pinching motion to capture the GQD. Together, these binding modes may represent early intermediates of the TA disaggregation mechanism. We find that functional groups capable of possessing a negative charge (*e.g.*, *COO*^-^, *O*^-^, and *S*^-^) produce impressive binding affinities. We propose that enriching these functionalizations during GQD synthesis and preparation, particularly sulfur as it is less studied, may yield more potent TA inhibitors and generalize to other amyloid pathologies with positively charged fibril cores.

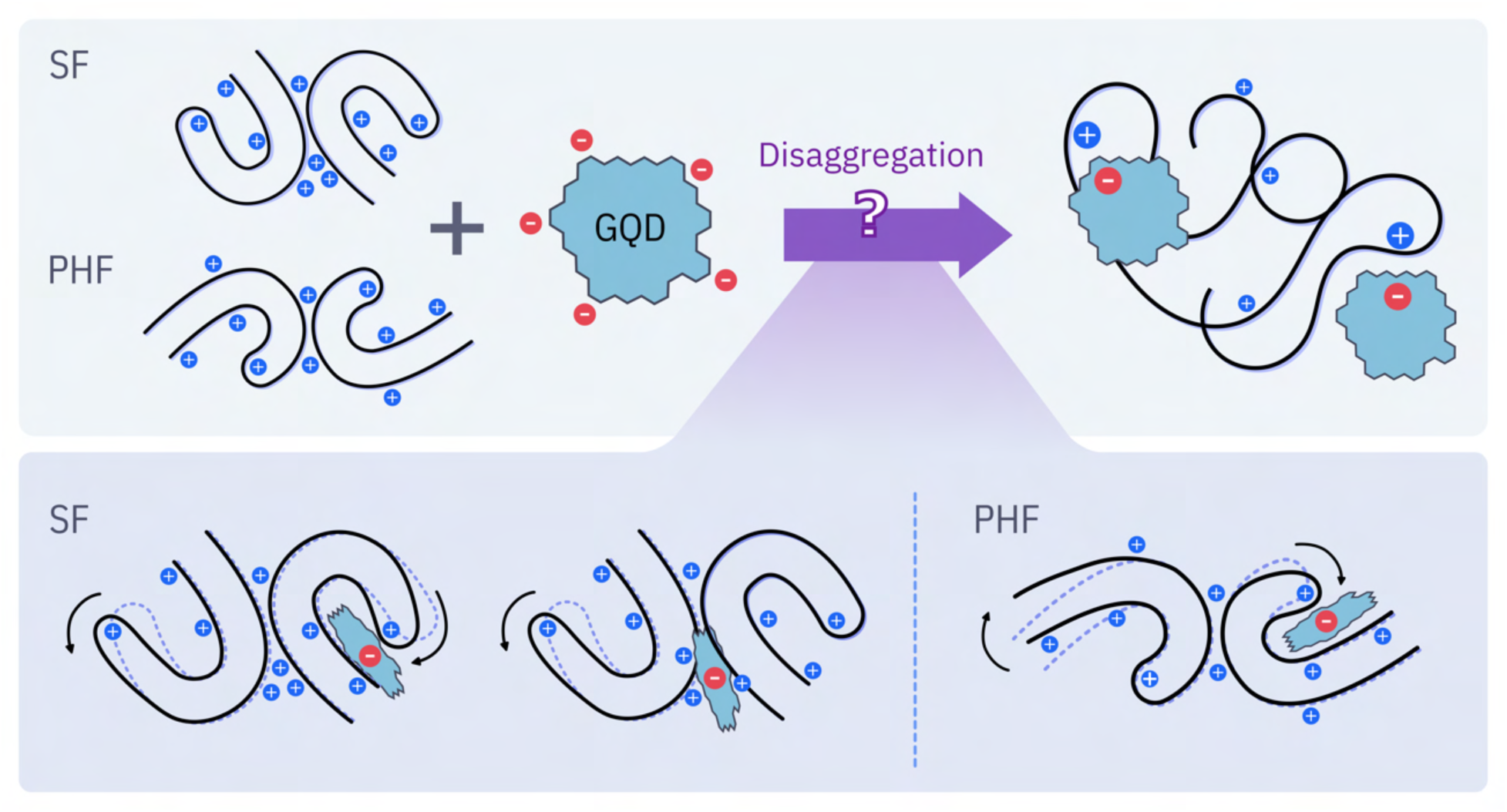

## 1 Introduction

Alzheimer’s disease (AD) is a widespread neurodegenerative disorder that has remained highly resistant to therapeutic interventions. It is projected that by 2050 there will be over 150 million AD cases globally [1], with the economic toll of AD and other dementias between 2020 and 2050 estimated at 14.5 trillion international dollars [2]. This underscores the urgent need to identify effective AD therapeutics, potentially through unconventional approaches. To date, there are three United States Food and Drug Administration-approved disease-modifying therapeutics for AD [3]: (1) Aducanumab (withdrawn); (2) Lecanemab; and (3) Donanemab. All of which are monoclonal antibodies that target amyloid-*β* (A*β*) in its soluble oligomeric or fibrillar forms and show modest clinical benefit. While amyloid-*β* and its aggregates have long been targeted in the amyloid-cascade hypothesis, mounting evidence indicates that Tau protein aggregates (TAs) correlate more strongly with cognitive decline [4–6]. Moreover, the removal of TAs improved cognitive functioning in AD [7, 8] and can even improve A*β* pathology [8]. Similar effects are noted in Tau-knockout models where A*β* toxicity is substantially reduced [9]. Hence, TAs represent a promising target for AD therapeutics and diagnostics.

Tau plays multiple physiological roles, most notably promoting microtubule assembly by stabilizing tubulin polymers. Tau exists as six main isoforms in the brain, differing by the number of N-terminal domains (0N, 1N, and 2N) and microtubule binding domains (3R and 4R). The 4R isoforms of Tau include a second fibril-forming hexapeptide, _275_VQIINK_280_ (numbering based on the 2N4R isoform), which is a stronger driver of aggregation than the _306_VQIVYK_311_ hexapeptide present in all six isoforms [10, 11]. In AD, TAs appear as two fibril polymorphs, straight filaments (SFs) and paired helical filaments (PHFs), with the latter being the predominant form [12]. In both polymorphs, the fibril core comprises sections of the third and fourth microtubule binding domains and is surrounded by a disordered fuzzy coat [13, 14]. For a therapeutic or diagnostic molecule to readily access the positively charged fibril core, it should be capable of penetrating the negatively charged fuzzy coat.

Being recognized by the 2023 Nobel Prize, quantum dots (QDs) garnered significant attention across many disciplines. Graphene QDs (GQDs) are of particular interest for therapeutic applications due to their low toxicity [15] and minimal effect on cell viability [16], ability to cross the blood-brain barrier, high solubility, and significant potential for functionalization [17]. GQDs have shown the ability to either prevent the aggregation or disassemble aggregates of: A*β* in AD [18, 19], *ε*-synuclein in Parkinson’s disease [20–22], human islet amyloid polypeptide in type-2 diabetes [23–25], TAR DNA-binding protein 43 which is implicated in amyotrophic lateral sclerosis (ALS), frontotemporal dementia, and AD [26, 27], fused in sarcoma in ALS [28], and even Tau in AD [29, 30]. The two aforementioned Tau experimental investigations featured approximately 8 nm diameter GQDs with carboxylate, ethylenediamine, D-cysteine, or D-cysteine alongside the non-covalent mxyl-NAP2 peptide functionalizations. By scanning different charge states, Zhu *et al.* [30] found that anionic GQDs, containing carboxylate groups, were effective at inhibiting Tau aggregation and reasonably effective at disassembling TAs, though less capable of penetrating Tau’s negatively charged fuzzy coat. To improve GQD selectivity, mxyl-NAP2, a TA capping *β*-bracelet peptide, was adsorbed onto the surface of a D-cysteine functionalized GQD. By saturating the GQD’s surface with the peptide, selectivity for inhibiting Tau over bovine serum albumin aggregation was markedly improved.

While Zhu *et al.* demonstrate impressive outcomes with their GQDs, the scope of tested functionalizations and charge states is limited. Furthermore, mechanistic insight into how GQDs interact with Tau at the atomic level remains to be elucidated. To better understand and generate GQDs that can optimally bind to and disrupt TAs, we must begin by identifying what covalent GQD functionalizations improve affinity to TAs. To accomplish this, we identified the putative GQD binding sites in brain-derived SF and PHF TA structures. We then performed exhaustive docking at each of these sites with our custom library of functionalized and variably charged GQDs. Finally, we evaluated the stability of the docked poses in large-scale molecular dynamics (MD) simulations. In each stage of the study, we compare results to the pristine (unfunctionalized) GQD to determine the effect of functionalization. This study aims to serve as a guide for future GQD development for AD and provide atomic-level insight into how GQDs interact with TAs in AD.

## 2 Methods

### 2.1 GQD Preparation and Parametrization

Functionalized GQDs were built based on our previously optimized pristine GQD structure [31]. Molecular geometries were first subjected to CREST-CENSO calculations [32, 33] to optimize the flexible functionalizations. Next, each GQD’s most populated conformer was selected and optimized at the HF/6-31G* level of theory with Gaussian 16 software [34]. A frequency calculation at the same level of theory was performed to confirm that the optimized geometries correspond to minima. CHELPG charges were derived according to the RESP protocol [35] and fit to the molecular mechanics grid with Antechamber [36], with custom force field parameters generated as needed with parmchk2.

### 2.2 Binding Site Identification

We determined the binding sites within our protein structures using a consensus among CavityPlus [37], FTMap [38], and P2Rank [39] web servers. Within each web server, the protein PDB IDs were inputted and default settings were used. We also considered pristine GQD binding sites that our group has previously identified through exhaustive docking [31] to multiple SF and PHF structures. We analyzed binding site electrostatics with the Adaptive Poisson-Boltzmann Solver algorithm (APBS, isovalue = 5) [40] in PyMOL [41] and hydrophobicity using the built-in molecular lipophilicity potential tool in ChimeraX 1.6 [42].

### 2.3 Protein Structure Preparation

The SF (PDB ID: 5O3T [13]) and PHF (PDB ID: 5O3L [13]) structures were retrieved the Protein Data Bank [43]. We selected these two TA structures because they had previously been demonstrated to behave well in MD simulations (unlike structures with fewer chains, as noted in [31]) and to allow for a direct comparison with our previous work. Protonation states were assigned using the H++ web server[44]. In preparation for docking, the fibril structures were placed in a cubic box with a 12 Å buffer of explicit OPC water [45] and 0.15 M NaCl with AMBER’s tLeap [46]. The ff99SB force field [47] was used for this step. The solvent and ions were first minimized using the steepest descent (SD) algorithm, then the protein was minimized using the SD algorithm, followed by the conjugate gradient algorithm. The minimized structure for each fibril was used for docking calculations. We performed normal mode analysis (NMA) on the SF and PHF crystal structures using the ANM 2.1 web server [48] with default settings. We visually analyzed normal modes using PyMOL [41].

### 2.4 Docking GQDs to SF and PHF Fibrils

For all docking simulations, the GQDs’ non-polar hydrogens were merged with their bonded heteroatom in AutoDockTools4 [49]. The minimized protein structures were processed by setting atom types to AD4, adding Kollman charges, and merging the non-polar hydrogens with their bonded heteroatom. We generated a cubic grid box of 66 *points*^3^ with 0.375 Å grid spacing with AutoGrid4, ensuring that each grid box axis was at least three times the largest GQD’s radius of gyration. We docked the GQDs to each binding site using AutoDOCK-GPU [50] with the Lamarckian Genetic Algorithm (LGA) for global optimization across 75 independent runs. Each run had a population of 200 and was allowed up to 5,000,000 energy evaluations and 50,000 generations. Local search was performed using the Pseudo-Solis and Wets algorithm with up to 500 iterations per pose.

To select GQD poses for MD simulations, the best-scoring pose for each GQD at each site was used. For each site, docking scores were normalized using Z-scores, calculated using the difference between a functionalized GQD’s docking score and that of the pristine GQD, divided by the standard deviation of docking scores within its respective binding site. Due to computational cost, no more than three functionalized GQDs were selected per site, and each GQD could be assigned to at most two of its best-scoring sites to maintain GQD diversity. GQDs were prioritized for site assignment based on their average Z-score for each TA. The best-scoring pristine GQD pose was included as a control at all sites. Docked interactions were identified with PLIP [51], except for *S*^-^ interactions, which were added manually. Prior to MD simulations, the docked heavy atom coordinates were mapped onto the RESP-generated GQD structure.

### 2.5 Molecular Dynamics (MD) Simulations of Selected Poses

MD systems were prepared using tLeap in AMBER23 [46] with the GAFF2 ligand force field and ff19SB protein force field [52]. Each system was solvated with OPC water molecules [45] in an octahedral box with a buffer of 12 Å between the outermost residue and the edge of the box. *Na^+^* and *Cl*^-^ counter ions were then used to neutralize and ionize each system to 0.15 M NaCl based on the number of water molecules [53]. All minimization and equilibration simulations were performed by MDProtocol [54], an automated in-house Python package using OpenMM 8.1.1 [55]. This package was used for two main reasons: (1) it is fully customizable and allows for independent treatment of position restraints across different atom groups; and (2) it supports the fitting of restraint weights according to any user-defined function. The second feature is especially important, as it prevents sharp energy spikes caused by abrupt adjustments in restraint weight (RW), which can greatly destabilize the system. We split each system into four groups of atoms with different initial (RWs): (i) protein backbone (PBB: N, CA, C, and O atoms, RW=10 kcalmol^-1^Å^2^); (ii) protein side-chains (PSC, RW=5 kcalmol^-1^Å^2^); (iii) ligand anchor atoms (LA, all GQD lattice carbons, RW=10 kcalmol^-1^Å^2^); and (iv) ligand "extensions" (LEX, all non-hydrogen, non lattice-carbon GQD atoms, RW=5 kcalmol^-1^Å^2^). We define the RWs using a decay function of the form

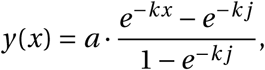

where a is the initial RW, and j is the step at which the RW becomes 0, and k determines the rate of decay, as calculated based on user input (*k*_0_):

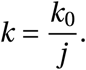

We performed our entire equilibration protocol in triplicate, starting from the same solvated and ionized complex. The first step of the protocol is to minimize the system until convergence in three stages with the Limited-memory Broyden–Fletcher–Goldfarb–Shanno algorithm: (1) minimize with all of the aforementioned RWs (Figure S1A); (2) minimize with only PBB and LA groups restraints (Figure S1B); and (3) minimize with no restraints (Figure S1C). Next, the fully restrained system is slowly heated from 100 to 300 K over 3 ns with the Nose-Hoover thermostat [56, 57] and H-bond lengths constrained, followed by an NVT equilibration at 300 K for 2 ns (Figure S1D,E). The Monte Carlo barostat [58] is then turned on and the RWs are gradually reduced over 15 ns. Specifically, the first 5 ns of NPT equilibration have the PSC and LEX restraints decreasing at a decay rate of 5 step^-1^ (Figure S1F,G). The following 10 ns have the PBB and LA restraints decreasing at decay rates of 10 and 5 step^-1^, respectively (Figure S1H,I). Finally, the system is simulated at NPT unrestrained for 5 ns (Figure S1D,E).

To transition this system to running with AMBER23 [59], we performed an additional 5 ns NPT equilibration with the Bussi thermostat [60] and H-bond distances constrained by SHAKE [61]. All production simulations were run in triplicate (333.33 ns each), yielding 1 *µ*s of data for each system.

### 2.6 Analysis of MD Simulations

Equilibration trajectories were processed using MDTraj [62] (automated by MDProtocol [54]) and production trajectories were processed with CPPTRAJ [63]. Visual trajectory analysis and figure generation were performed using ChimeraX 1.6 software [42]. We assessed global protein conformation changes with root-mean square deviation (RMSD) calculations and residue-level fluctuations with RMS fluctuation (RMSF) of the aligned protein backbone atoms. Fibril secondary structure changes were monitored with the DSSP algorithm [64]. Radius of gyration (*R_g_*) calculations tracked the compactness of each protofilament. In addition, we calculated the distance between ARG349 and LYS375 in the central chain of each protofilament to measure the degree of conformational pinching present. To determine whether the GQDs impact the buried surface area at the protofilament interface, we calculate the buried surface area (BSA) of interface residues by taking the difference between the SASA for the protofilament in the presence and absence of the other protofilament. We combined the above replicate data for each system and calculated the degree of overlap with the respective apo systems’ densities using a common number of bins, allowing for global nonparametric comparisons. More specifically, for each statistic (RMSD, BSA, *R_g_*, and ARG349-LYS375 distance) and for each TA, we determined the number of bins using the Freedman–Diaconis rule [65]. In addition, we performed a Dynamic Cross Correlation (DCC) analysis [66] on the aligned alpha carbon coordinates of each protofilament’s central chain.

To obtain representative structures of the apo systems, we first performed Principal Component Analysis (PCA) on the combined alpha carbon coordinates (aligned to a common reference structure) for each apo system using the scikit-learn package [67]. We retained as many PC vectors as needed to account for >95 % of variance for subsequent agglomerative hierarchical clustering with ward linkage. Based on the largest Silhouette score value, the number of clusters was chosen for each system [68]. Within each cluster, the most representative structure was extracted. The free energy surface (FES) was constructed with respect to the first two PCs, with the most representative structures labeled.

We quantify the binding free energies (BFEs) of the GQDs to the TAs using 200 frames from the last 50 ns of production simulations with the molecular mechanics Poisson-Boltzmann surface area (MM/PBSA) method. We set the internal dielectric set to 4 due to the degree of charge and exposure of our binding sites [69] and the ionic strength to 0.15 M. We decompose the BFEs into electrostatic and Van der Waals contributions, from which we analyze the top residue contributions to the BFEs. While MM/PBSA is generally unreliable for absolute BFE predictions, especially without empirical correction, it remains a useful screening tool for determining the relative BFEs of a series of structurally related ligands, such as our GQDs [70]. We also calculate the BSA for the GQD when bound to the TAs and in isolation.

## 3 Results and Discussion

### 3.1 Binding Site Identification

GQDs are expected to be capable of binding to TAs through diverse binding modes. We utilized three independent software tools to predict consensus binding sites in both TAs: CavityPlus [37], FTMap [38], and P2Rank [39]. Nevertheless, these software tools were not designed to predict binding sites for such unconventionally large ligands, hence we also consider the binding sites that we previously identified by exhaustively docking a pristine GQD to TAs [31].

Analysis of the SF structure reveals that the consensus among the three programs identifies the C-shaped curve and the small protofilament interface as viable binding sites. In our previous study of pristine GQDs [31], we identified the C-shaped curve binding mode, along with the large protofilament interface and the fibril termini (a surface interaction containing the PHF6 region), but not the small protofilament interface. We decided to pursue docking simulations at the C-shaped curve, at both protofilament interfaces, and at the fibril termini. The C-shaped curve binding site contains a mix of residue types and has a primarily positive electrostatic surface (Figure S2). This binding site is reasonably narrow, and a 2 nm diameter GQDs barely fits when interacting facially with ILE360. The small protofilament interface is mostly polar, with a positive electrostatic surface but limited depth for GQD binding within its groove. The large protofilament interface possesses a deep groove, whose opposing faces are polar and hydrophobic and exhibit a highly positive electrostatic surface. The termini binding site has a mixed residue composition with a mildly positive electrostatic surface and is an important target since it contains the pathogenic PHF6 region, where GQD binding may cap TA elongation.

In the PHF structure, the consensus among the three programs indicates that there is a viable binding site at the protofilament interface. We previously identified this binding site, along with the C-shaped curve and fibril termini [31]; therefore, we also considered these regions in our docking simulations. Unlike the SF, the two PHF protofilament interface sites are symmetric; each is polar, shallow, features a small hydrophobic patch, and lacks a highly positive electrostatic surface (Figure S3). The PHF C-shaped curve and termini binding sites are nearly identical to the SFs’; however, the former is slightly narrower. Having established the relevant binding sites in both TAs, we next evaluate GQD interactions through docking simulations.

### 3.2 Docking GQDs to Tau Aggregates

We performed docking with AutoDOCK-GPU [50] with a large amount of sampling to ensure the optimal placement of both the GQD lattice and its flexible edge groups. In each binding site, we collect each GQD’s best docking score, listed in Tables S1 and S2, with interactions identified by PLIP [51]. Figure 1A,B presents the placement of all best-scoring docked poses for each GQD, colored by docking score. Given the large positive net charge of Tau’s fibril core, the anionic GQDs produce much more negative (favorable) docking scores, followed by the neutral and positive GQDs (Figures 1C,D, S4, and S5). Representative docked poses of pristine and an anionic GQD are provided in Figure 2. The docking scores range from -29.35 to 7.19 kcalmol^-1^ in SFs, and from -17.67 to 5.89 kcalmol^-1^ in PHFs. This difference results from SFs possessing the large protofilament interface with a high degree of positive charge, compared to the more charge-balanced protofilament interfaces in PHFs. In SFs, the anionic GQDs have an apparent preference for binding to the large protofilament interface, whereas the neutral and positive GQDs have no clear site preference. In PHFs, GQDs generally favor binding to the C-shaped curve and termini rather than the protofilament interface; however, there is no distinctly preferred site. Nevertheless, in the apparent absence of a dominant binding mode (except for anionic GQDs in SFs), these widespread high-affinity interactions with TAs suggest that non-specific binding may be the driving force leading to experimental disaggregation [30], which we further investigate using MD simulations.

**Figure 1:**
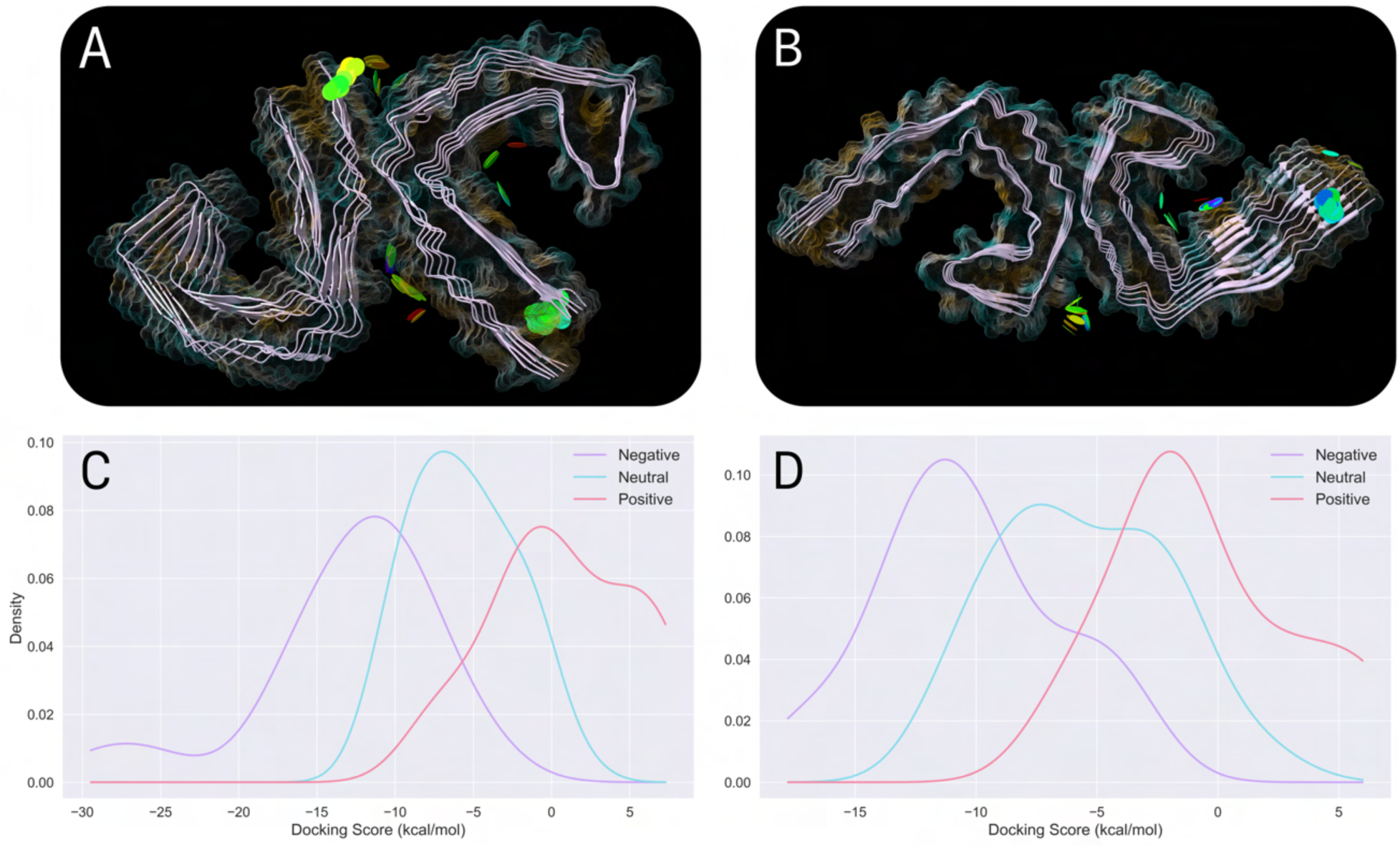
Summary of docking study: Best-scoring pose for each GQD, represented as discs and colored by docking score (most negative (violet) to most positive (red) using a reversed rainbow scale), bound to SF (A) or PHF (B) and colored by hydrophobicity in ChimeraX 1.6 [42]; Kernel density estimate (KDE) of GQD docking scores with SF (C) or PHF (D), separated by charge state.

**Figure 2:**
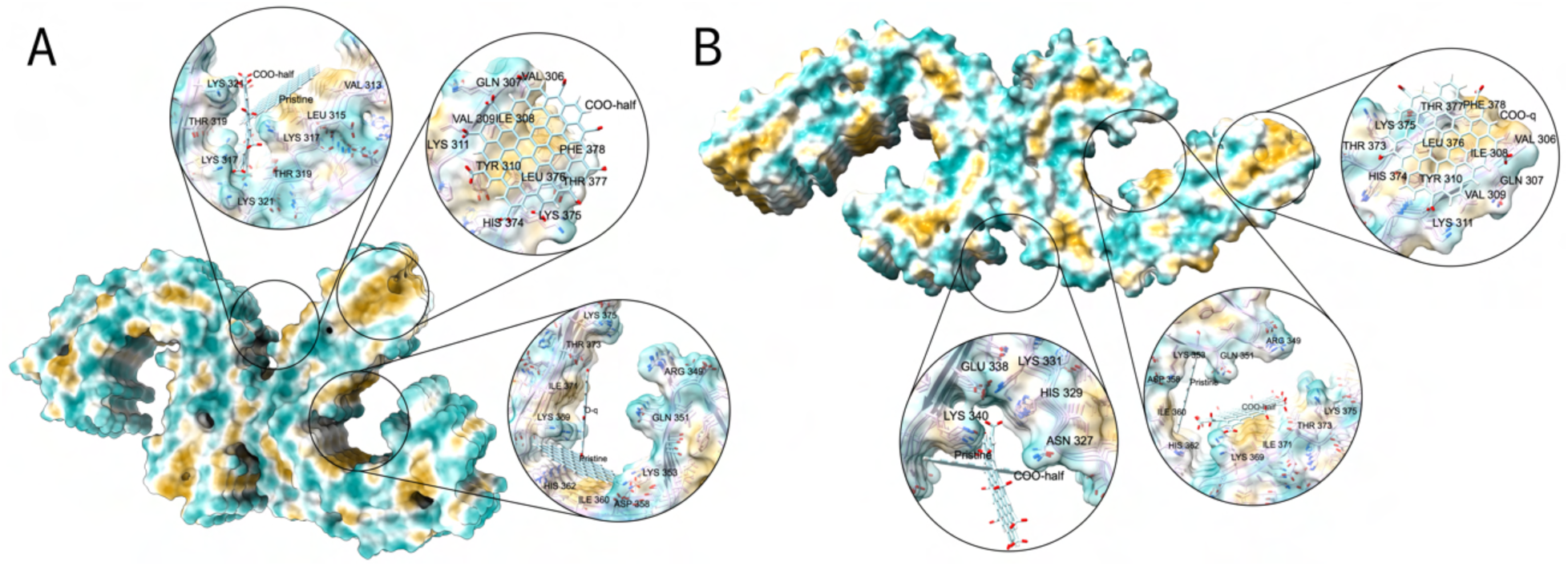
Representative docked poses for pristine and an anionic GQD in SF (A) and PHF (B). Binding at the SF small protofilament interface is not shown due to its unfavorability. The protein surface is colored by its hydrophobicity, calculated by ChimeraX 1.6 [42].

In the SF’s C-shaped curve, the pristine GQD docked similarly to our previous study, though it interacts more facially with ILE360 while also forming contacts with ASP358 and LYS369 (Figure 2A). Both ASP358 and ILE360 are important residues for disease-specific binding of TAs, along with GLN351 and LYS353, which appear to be nearly forming an interaction. On the other hand, in PHF’s C-shaped curve, the pristine GQD binds nearly identically to what was observed in our previous work (Figure 2B) [31], interacting most with ILE360 and LYS353, while also forming a weak interaction with GLN351. Upon functionalization, the GQDs consistently occupy two distinct binding modes in both TAs: (1) interacting with the same hydrophobic patch as pristine in SFs, though gaining new interactions with nearby polar and charged residues; or (2) interacting with a neighboring hydrophobic surface that is nestled between two positively charged patches, typically interacting with LYS369, ILE371, and THR373. The former mode is observed for three-quarters (half) of the functionalized GQDs in SFs (PHFs) that lack a high degree of negative charge (*e.g.*, half functionalized *COO*^-^, *O*^-^, and *S*^-^). The latter mode is occupied mainly by highly anionic GQDs and produces the most negative docking scores in this binding site.

In the termini binding sites, the pristine GQD interacts with both the PHF6 region and its opposing chain via a surface interaction (Figure 2). Almost all functionalized GQDs dock favorably in the same manner as pristine, indicating that this interaction represents a consistent binding mode across GQD functionalizations and is not governed exclusively by electrostatics.

Docking attempts to the SF small protofilament interface rarely position the GQDs within the groove. Instead, the GQDs tend to bind to the fibril termini in the same way as the termini binding site. This is consistent with our previous work [31], which indicated that the pristine GQD does not bind favorably to this interface, whereas it does to the nearby PHF6 region. Six of the functionalized GQDs are capable of burying into the interface groove and achieving more negative docking scores. These were almost all anionic GQDs, as expected given this site’s positive electrostatic surface. An additional binding mode that bridges both protofilaments is observed but is generally unfavorable. In the large SF protofilament interface, the pristine GQD interacts facially with a hydrophobic patch primarily composed of LEU315 and does not bury deeply into the groove, consistent with our previous study. The neutral GQDs, especially those incapable of hydrogen bonding, bind similarly to pristine. The anionic GQDs and neutral GQDs capable of hydrogen bonding insert more deeply into the groove and interact with LYS317, THR319, and LYS321 (see, for example Figure 2A). Anionic GQDs in this binding pose achieved highly negative (favourable) docking scores relative to the other binding sites, indicating that this is likely their dominant binding site in SFs.

In the PHF protofilament interface, the pristine GQD bridges both protofilaments, interacting with LYS340 and GLU342, but does not bury into the interface groove (Figure 2B). Nevertheless, this binding groove is shallow and provides little volume for GQDs to occupy. The majority of the functionalized GQDs bind weakly in a mode similar to pristine, with the anionic GQDs achieving the most negative docking scores. Four of the functionalized GQDs somewhat interact within the groove but do not achieve appreciably negative docking scores. Overall, this binding site is less favorable than the SF’s large protofilament interface.

Altogether, in TAs, the neutral and quarter functionalized GQDs generally adopt a binding mode distinct from that of the anionic and half functionalized GQDs. The anionic GQDs tend to produce much more negative docking scores than all other GQDs, which aligns with TAs being highly positively charged. The cationic GQDs produce near-zero or positive docking scores, suggesting that they are unsuitable for TA binding. These results are in line with experimental work showing that anionic GQDs generally exhibit enhanced ability to disaggregate Tau, relative to cationic GQDs [30].

#### 3.2.1 Determining the Influence of Charged State on Docking Score

Some of the GQDs were designed to have different charge states using the same functional group. To circumvent the computational complexity of identifying their exact distribution of charged states, we instead use model systems of purely neutral or charged functionalizations (*e.g.*, we have all COOH groups in one GQD and all *COO*^-^ groups in its charged counterpart). We treat these models as the extremes of protonation states that enclose plausible behavior within the variable binding sites microenvironments; moreover, we assume that more physical mixed protonation states will score between these extremes. When switching from a neutral GQD to its anionic counterpart in the same binding site, the docking scores decrease (became more favorable) on average by 11.81 and 5.41 kcalmol^-1^ for half and quarter functionalizations, respectively (Tables S1 and S2). On the other hand, when switching from a neutral GQD to its cationic counterpart in the same binding site (*N H*_2_ to *^+^N H*_3_), the docking scores increase (became less favorable) on average by 5.54 and 2.31 kcalmol^-1^ for half and quarter functionalizations, respectively. Overall, half functionalized GQDs tend to at least double the docking score impact of their quarter functionalized counterparts (Figure S6), indicating a strong sensitivity to charge density. These trends, however, should not be directly extrapolated to the behavior of different heteroatoms incorporated within the lattice (*e.g.*, B, N, S), which are expected to have a more modest effect on the binding affinity. Instead, lattice functionalization can be better suited for tuning the optical properties (*e.g.*, absorption range) of GQDs [71], an area currently under investigation for developing GQD-based diagnostics and theranostics.

#### 3.2.2 Selecting GQDs for MD Simulations

We selected GQDs for MD simulations based on the extent of normalized binding affinity improvement, relative to the pristine GQD, obtained at each site while maintaining GQD diversity (details provided in Methods). All selected GQDs are anionic (Figures S7, S8, and Table S3), in agreement with experiments which show anionic GQDs effectively inhibit Tau aggregation and induce disaggregation [30]. To gauge the effect of functionalization, we took the pristine GQD’s best-scoring pose in each binding site for MD simulations. There is a general mix between quarter and half functionalized GQDs at each site, except SF’s large protofilament interface. While docking offers an effective overview of GQD binding, it cannot reveal the persistence or stability of these interactions, nor their effect on TA stability. To address this, we next performed MD simulations on the selected GQDs.

### 3.3 Molecular Dynamics Simulations of Docked Poses

#### 3.3.1 Understanding the Structural Dynamics of Tau Aggregates

Studying protein structural dynamics can reveal how collective motions induce transient conformations, some of which can be exploited for ligand binding to stabilize a specific conformational state. We first analyze the MD simulations of SF and PHF apo systems in triplicate to establish a baseline for comparison with the GQD-TA systems and to understand how GQD binding affects protein structural dynamics.

Apo SF MD replicates produce similar results in general, with an RMSD relative to the minimized crystal structure of 4.22 ± 0.88 Å overall (Table 1). From RMSF analysis, it is evident that the largest structural changes occur at the terminal residues, which often slightly unpeel from the protofilament. There is no appreciable loss in secondary structure, altogether implying that SFs are stable in our simulations. The protofilament *R_g_* values stabilize at similar levels, with Protofilament 1 being somewhat more compact than Protofilament 2 throughout the simulations (24.65 ± 0.43 versus 25.26 ± 0.69 Å, respectively). The *R_g_* values tend to oscillate due to a collective motion in which each protofilament undergoes a constriction resembling a pinching motion followed by an opposite, outward unpinching motion. By quantifying the BSA between the protofilaments and monitoring the distance between ARG349 and LYS375 on both faces of the pinching interface (Figures 3A,B and S9), we observe that protofilament pinching and unpinching have a minimal effect on BSA. Often, when one protofilament pinches, the other exhibits a compensatory motion that stabilizes the BSA, suggesting a possible mechanism for maintaining protofilament interface stability.

**Figure 3:**
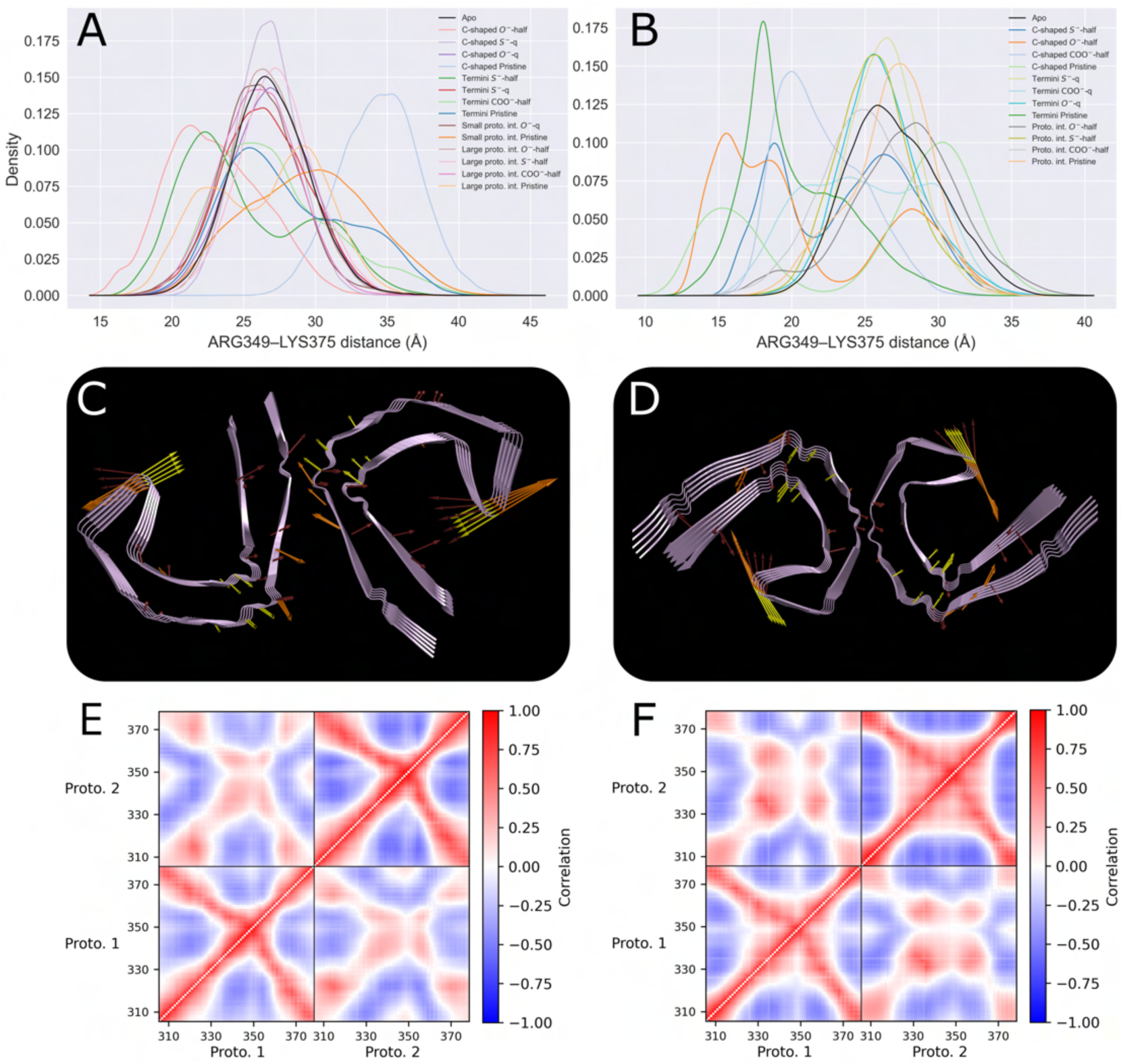
Analysis of the pinching conformation in TAs: KDE of ARG349-LYS375 distances for Protofilament 1 in SFs (A) and PHFs (B); Three representative low-lying, in-plane normal modes of symmetric and asymmetric apo pinching conformations (mode amplitudes increased for visualization) for SFs (C) and PHFs (D); DCC plot of apo inner protofilament chains for SFs (E) and PHFs (F).

**Table 1:**
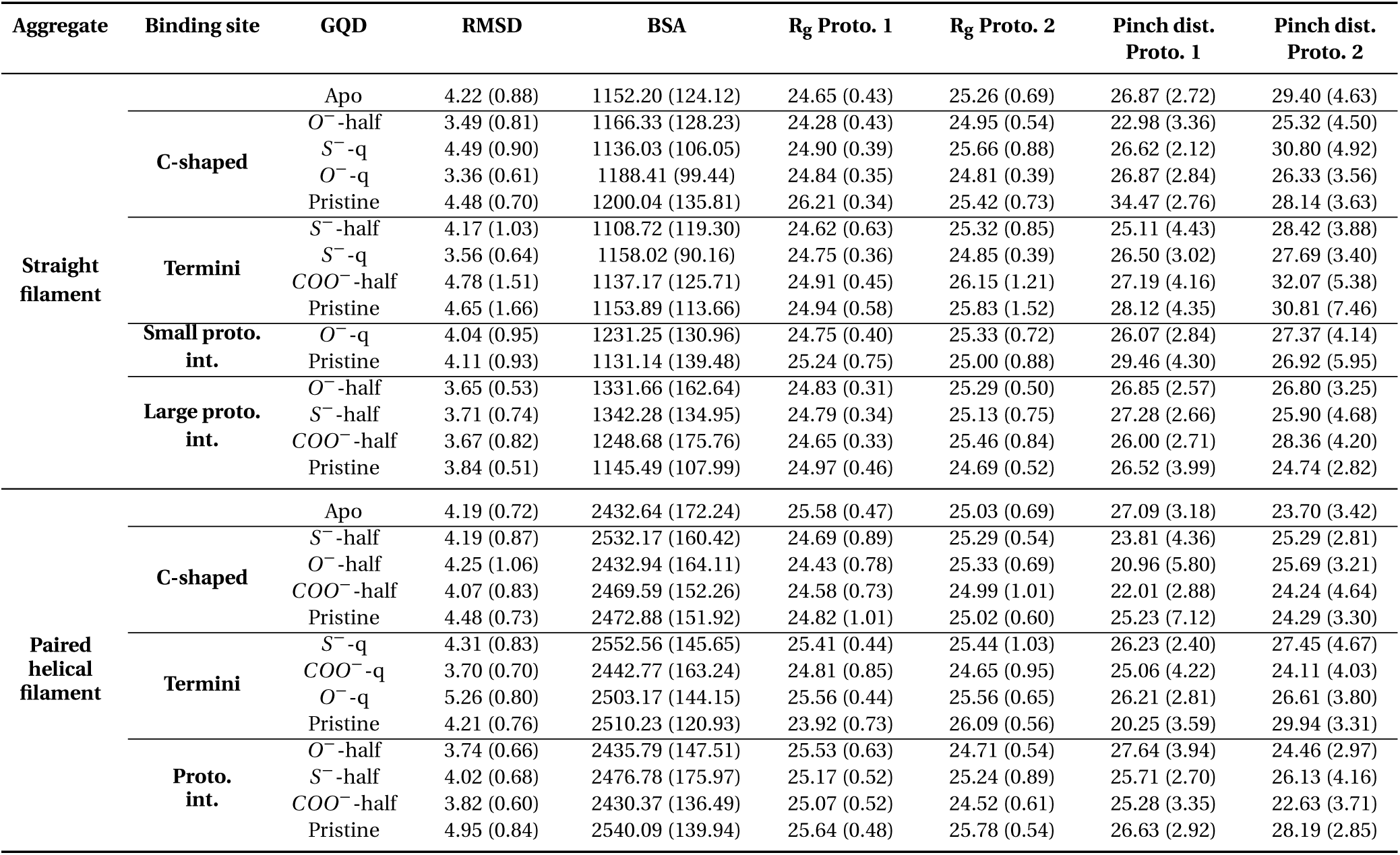
Summary of MD production simulation statistics for SF and PHF. All values are presented as the pooled mean (standard deviation) across all replicates in Å (or Å^2^ for BSA).

To further investigate this pinching motion, we performed NMA on the alpha carbons of the SF crystal structure. NMA reveals that five of the ten most energetically accessible vibrational modes involve this pinching motion in at least one of the protofilaments (see three representative modes in Figure 3C,D). From our MD DCC plots, we observe a negative correlation within-protofilament between the hydrophobic turn motif (residues 341–354) and termini regions which are involved in the pinching motion (Figure 3E), further supporting the existence of this collective motion. Between protofilaments, a positive correlation exists between the termini of one protofilament and the hydrophobic turn motif of the other, hinting that the pinching motion occurs asymmetrically. The electrostatic repulsion arising from the induced proximity of ARG349 and LYS375 in the pinched conformation likely regulates this collective motion and helps maintain TA stability. Finally, from clustering the coordinates of SF alpha carbons (details provided in Methods), we obtain two clusters whose most representative structures are quite similar in free energy, with the larger cluster (c0) corresponding to a slightly pinched conformation (Figure S10).

Apo PHF MD simulations behave similarly to the apo SF systems. They produce an RMSD of 4.19 ± 0.72 Å across replicates (Table 1), exhibit sporadic terminal motions, and show minimal loss in secondary structure. In this system, Protofilaments 1 and 2 now have more similar *R_g_* values (25.58 ± 0.47 versus 25.03 ± 0.69 Å respectively) but markedly different distances between ARG349 and LYS375 (27.09 ± 3.18 versus 23.70 ± 3.42 Å, Figures 3B and S11). Again, we notice that the pinching motion has a minimal effect on BSA and is often compensated by the other protofilament unpinching. NMA uncovers that five of the ten lowest-lying vibrational modes involve the pinching motion (see three representative modes in Figure 3D). This pinching motion is corroborated in the DCC plots, which exhibit the same trends as SF but with slightly stronger correlations (Figure 3F). We again identified two clusters whose most representative structures are energetically equivalent (Figure S12). Each structure contains one somewhat pinched and one somewhat unpinched protofilament, in agreement with our previous analyses. Together, this indicates that TAs readily undergo an energetically accessible pinching motion that may enable unique ligand binding modes, either via the increased residue solvent exposure in the protofilament interface grooves or the decreased volume in the C-shaped curve region.

#### 3.3.2 Investigating the Dynamics and Stability of GQD Binding: C-Shaped Curve

Simulations of pristine and functionalized GQDs within the SF and PHF C-shaped curves show protein stability comparable to the apo systems, with protein backbone RMSD values across replicates ranging from 3.36 to 4.49 Å (Table 1). These values represent overlaps of 0.57 to 0.88 and 0.80 to 0.91 with the SF and PHF apo RMSD density distributions, respectively (Table 2), where 1.00 indicates perfect overlap. SF beta sheet content tends to slightly increase, whereas the opposite trend is observed in PHF systems. Interestingly, the MM/PBSA BFE values are strongly correlated (*R*^2^=0.85) with the docking scores of the selected GQD poses across all binding sites (Figure S13). While this agreement suggests general consistency between the two approaches, it should be interpreted cautiously given the approximations inherent to both methods.

In SFs, the bound protofilament (Protofilament 1) shows smaller *R_g_* values than the unbound protofilament (Protofilament 2) for half functionalized and quarter functionalized *O*^-^ and *S*^-^, respectively, while quarter functionalized *O*^-^ yielded near-identical values. On the other hand, the *R_g_* value of Protofilament 1 in the pristine system is, on average, 0.79 Å larger than Protofilament 2 and 1.56 Å larger than the apo system. The *R_g_* value of Protofilament 1 represents an overlap of only 0.06 with the apo system, indicating that it adopts a distinct conformation. All half functionalized *O*^-^ GQD replicates remained locked in the pinched conformation, with the GQD extending from the hydrophobic patch near the C-shaped curve and bridging the opposite interface. In this system, the distance between ARG349 and LYS375 decreases by 3.89 and 4.08 Å relative to the apo system for Protofilaments 1 and 2, respectively (Table 2). This system yields low overlap in distance values with the apo system (0.51 and 0.64 for Protofilaments 1 and 2, respectively). In addition, the RMSD of this system decreases by 0.74 Å relative to the apo system, suggesting a global reduction in flexibility. The half functionalized *O*^-^ system produces BFE values between -80 and -71 kcalmol^-1^ (Table S4), with electrostatic interactions dominating, primarily involving ARG349, LYS369, LYS375, and occasionally LYS353 (Figure 4A).

**Figure 4:**
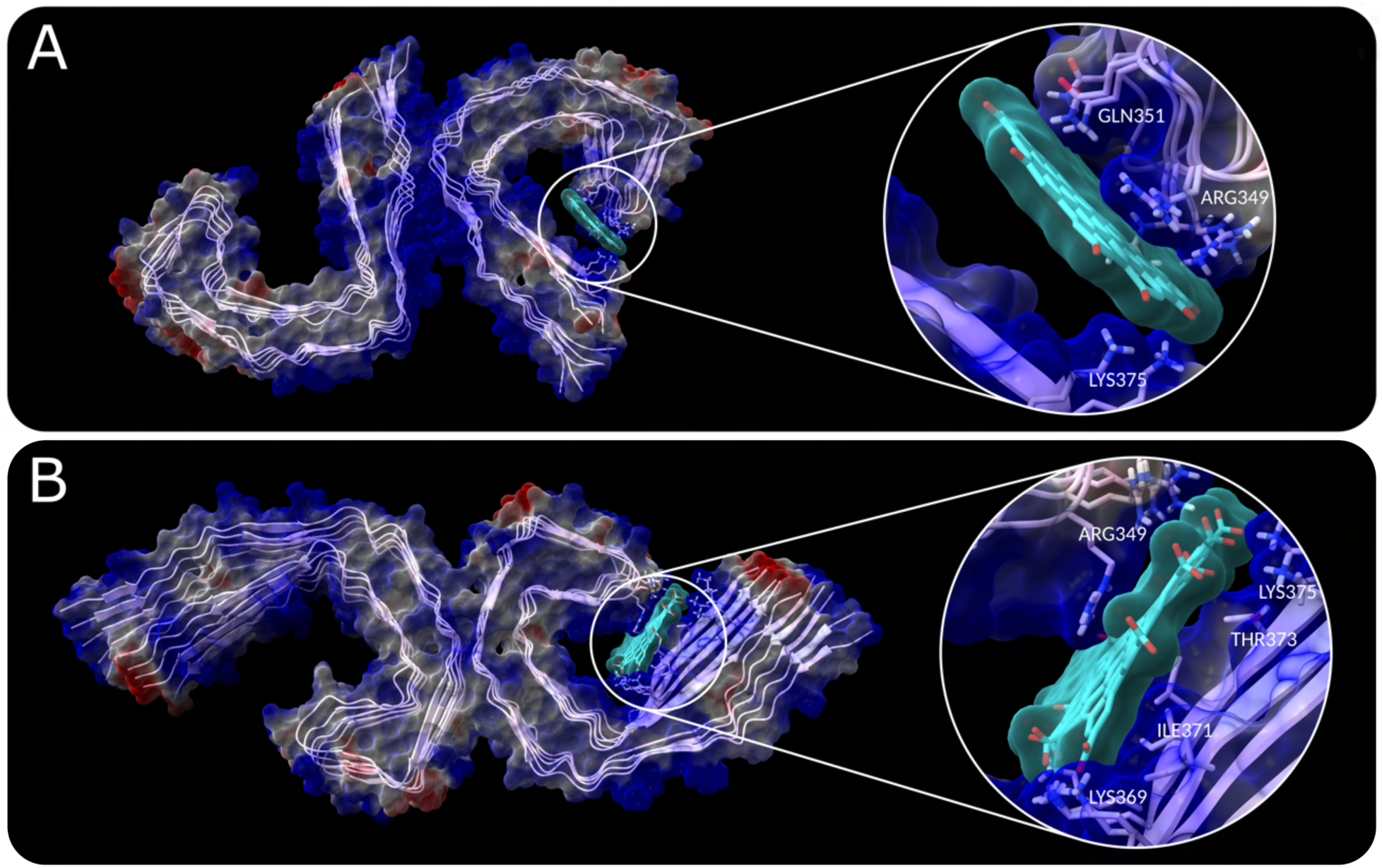
Most representative conformation from clustering of *O*^-^-half replicate 1 in SF (A) and *COO*^-^- half replicate 3 in PHF (B). The protein’s surface is colored based on its electrostatics, calculated by ChimeraX 1.6 [42] tial stages of GQD-TA interactions that lead to disaggregation [30]. In addition, we expect that larger GQDs than in the present study should be capable of forming this interaction. Nevertheless, the TA destabilization and disaggregation process can take several hours, as monitored with epigallocatechin gallate (EGCG) [10], a timescale inherently inaccessible to our simulations, which capture only the immediate effects of ligand binding.

**Table 2:**
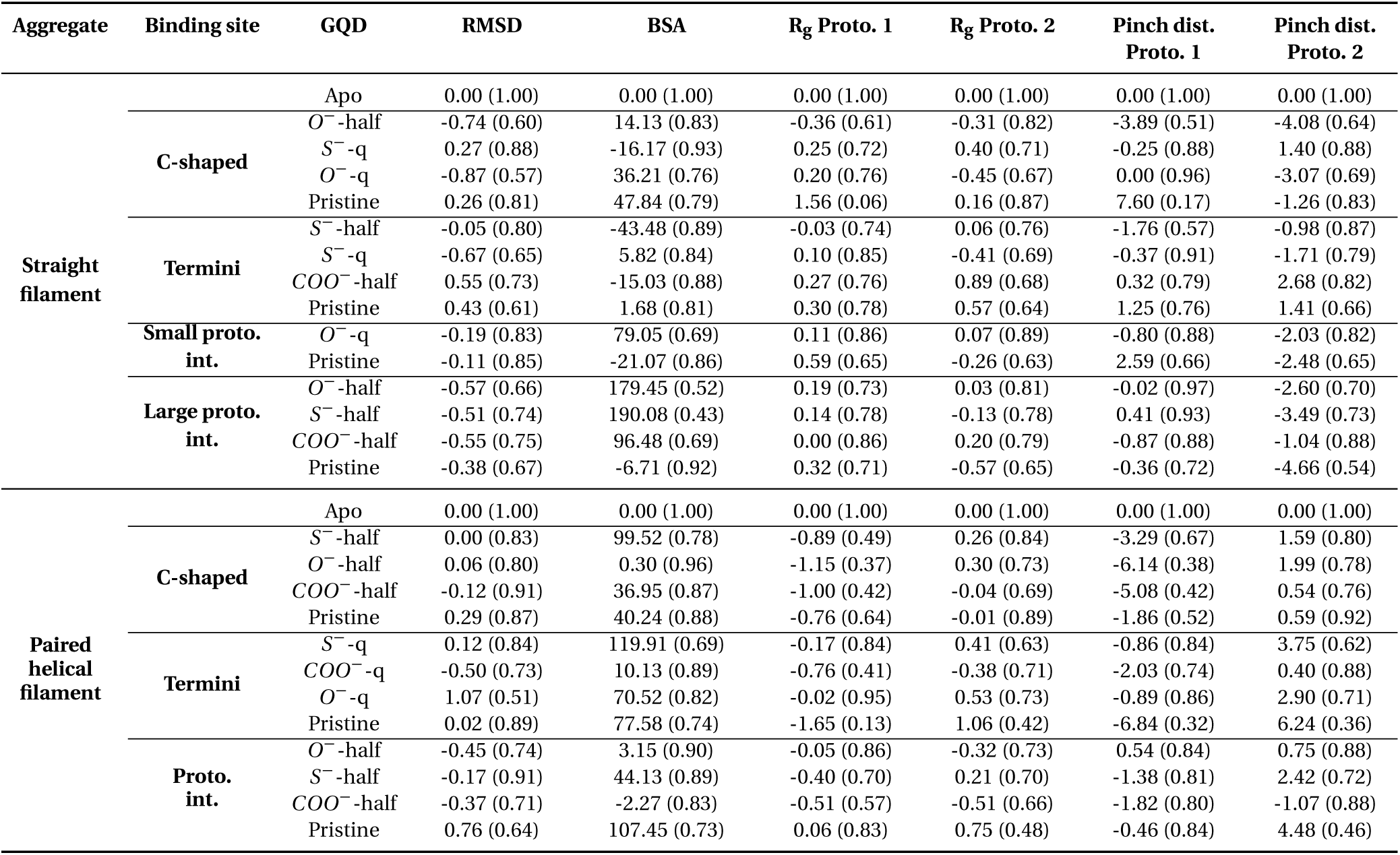
Summary of MD production simulation statistics relative to the apo SF and PHF systems. All values are presented as Δmean in Å (or Å^2^ for BSA) and (degree of overlap) relative to the apo system.

Interestingly, quarter functionalized *O*^-^ has the same starting pose as half functionalized *O*^-^ but does not form the bridged pinched state due to insufficient negative charge to form these electrostatic interactions. Quarter functionalized *O*^-^ interacts most with LYS369, ILE371, THR373, and LYS375 in the two replicates that remain in the docked pose, with BFE values between -54 and -51 kcalmol^-1^ (Table S4). The other replicate translated to the opposite surface of the pinching interaction (the hydrophobic turn motif), yielding slightly weaker binding (-46.48 ± 3.51 kcalmol^-1^). Quarter functionalized *S*^-^ and pristine bind similarly to each other in the C-shaped curve’s other hydrophobic surface, interacting with ILE360, HIS362, and PRO364, with the former also establishing electrostatic interactions with LYS353 and LYS369, yielding BFE values between -50 and -47 kcalmol^-1^ and around -18 kcalmol^-1^, respectively. Additionally, the quarter functionalized *S*^-^ and pristine systems show modest increases in RMSD relative to the apo system.

In PHFs, the pristine GQD maintains its docked pose, interacting with GLN351, LYS353, and ILE360, yielding BFE values between -26 and -20 kcalmol^-1^ (Table S4). In this system, the bound protofilament (Protofilament 1) *R_g_* values are, on average, 0.76 Å smaller than in the apo system (Table 2), while RMSD values are 0.29 Å larger than in the apo system. In PHF systems with half functionalized *S*^-^, *O*^-^, or *COO*^-^, the bound protofilament adopts the pinched conformation in eight of the nine simulations, with the GQD bridging the interface and forming the same interactions as the half functionalized *O*^-^ GQD in SFs (Figure 4B). Unlike with SFs, the PHF functionalized systems present minimal deviations in their RMSD values relative to apo. These systems yield overlaps of 0.67, 0.38, and 0.42 with the apo system’s bound protofilament ARG349 to LYS375 distances and have average distance reductions by 3.29, 6.14, and 5.08 Å, respectively (Table 2). Accordingly, the unbound protofilament’s ARG349 to LYS375 distances increased by 1.59, 1.99, and 0.54 Å relative to the apo system, respectively (Table 2 and Figure S11). These pinched poses correspond to the most negative BFE values in PHFs, ranging from -93 to -63 kcalmol^-1^, suggesting that this is the preferred binding mode for anionic GQDs in PHFs, in contrast to the docking study, which identified no binding mode preference. BFE values were especially negative for GQDs that formed extensive interaction interfaces, rather than primarily edge-based interactions.

The conformational asymmetry induced in the protofilament upon the binding of a highly functionalized anionic GQD may generate strain within SFs and PHFs and could hint at the ini-

#### 3.3.3 Investigating the Dynamics and Stability of GQD Binding: Termini

Among the 24 bound simulations, 23 remained in their docked pose with SFs and PHFs, forming a surface interaction with both terminal chains, centered marginally on the PHF6 region (Figure 5A,B), emphasizing the stability of this interaction. In the simulation deviating from this trend, the half functionalized *S*^-^ GQD migrated to become pinched by SF’s Protofilament 2, yielding a more favorable BFE than the other simulations. These complexes produce RMSDs between 3.56 and 4.78 Å (3.70 and 5.26 Å) in SF (PHF) (Table 1), with no consistent shift in stability across systems. SF *R_g_* values are typically slightly larger than the apo system in both protofilaments, whereas in PHFs, Protofilament 1 (2) has smaller (larger) *R_g_* values than the apo system. In SF, the pristine GQD achieves BFE values between -18 and -14 kcalmol^-1^, while the half functionalized *S*^-^ and *COO*^-^ and the quarter functionalized *S*^-^ yield BFE values from -56 to -53 (-72.92 ± 4.14 when pinched), -68 to -62, and -44 to -38 kcalmol^-1^, respectively (Table S4). In SFs, the functionalized GQDs interact most strongly with NVAL306, GLN307, VAL309, LYS311, THR377, and LEU376. The pristine GQD bound to the PHF achieves BFE values between -22 and -18 kcalmol^-1^, while the quarter functionalized *S*^-^, *COO*^-^, and *O*^-^ yield BFE values from -44 to -40, -41 to -40, and -43 to -40 kcalmol^-1^, respectively. In PHFs, the functionalized GQDs interact most strongly with NVAL306, ILE308, TYR310, LYS311, PRO312, LYS370, LYS375, and LEU376. This binding mode is favorable but not expected to be the dominant binding mode due to its modest BFE values relative to the other binding sites.

**Figure 5:**
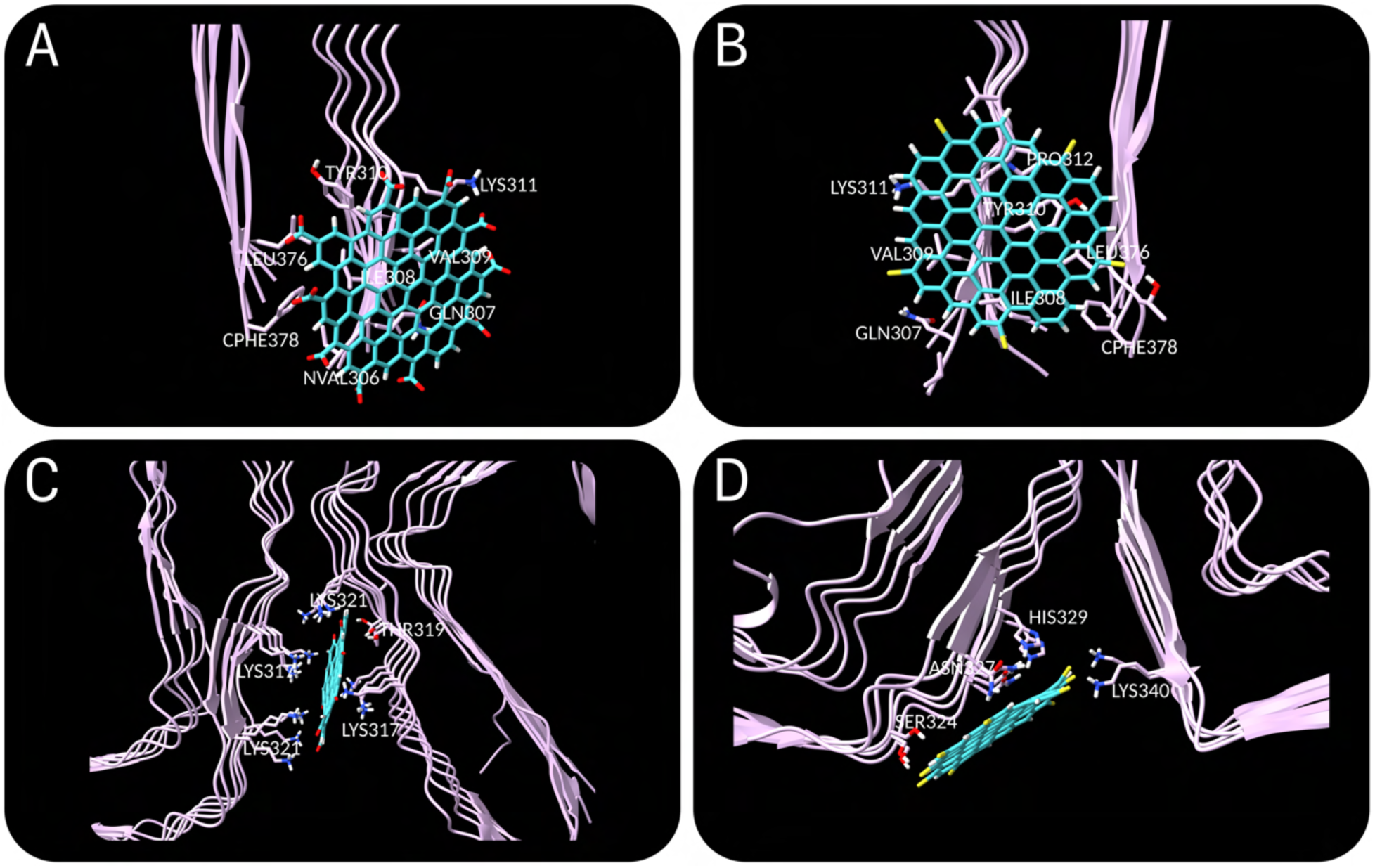
Snapshots of the most representative complexes for GQDs with highly negative BFEs at the SF termini (A, *COO*^-^-half replicate 2); PHF termini (B, *S*^-^-q replicate 3); SF large protofilament interface (C, *O*^-^-half replicate 3); and PHF protofilament interface (D, *S*^-^-half replicate 2).

#### 3.3.4 Investigating the Dynamics and Stability of GQD Binding: Protofilament Interfaces

Given the arrangement of the protofilaments, SFs have two unique protofilament interfaces whereas PHFs have only one (Figures S2 and S3). In agreement with our docking study and previous work [31], we confirm that the small interface in SFs is minimally receptive to GQD binding, regardless of the functionalization. Instead, the GQDs predominantly translate to form surface interactions with the adjacent PHF6 region (similar to the termini binding mode). This reduced propensity likely arises from the absence of a deep binding groove or hydrophobic patch at the small interface.

On the other hand, the large protofilament interface in SFs is primed for anionic GQD binding, given its high degree of positive charge and the presence of a hydrophobic patch (VAL313 and LEU315). In this binding site, the GQDs show minimal displacement and maintain their docked interactions during MD simulations, indicating stable binding. The pristine GQD retains its interactions with LEU315, VAL313, and LYS317, but also develops interactions with ASP314 and LYS321. One pristine GQD replicate slightly decreased its interaction depth within the groove, resulting in a less favorable BFE value of 0.44 ± 4.50 kcalmol^-1^ and GQD BSA of 160 Å^2^. Conversely, the other two pristine replicates have higher GQD BSAs (250 and 225 Å^2^), yielding slightly more negative BFE values of -5.20 ± 4.63 and -4.66 ± 5.03 kcalmol^-1^, respectively (Table S4). In agreement with our previous investigation, the pristine GQD is not expected to bind appreciably at this site.

The anionic GQDs (half functionalized *O*^-^, *S*^-^, and *COO*^-^) in this binding site achieve the most exaggerated BFE values in this study, ranging between -162 and -112 kcalmol^-1^ (Table S4) and have larger GQD BSA values than pristine. As previously noted, these BFE values should not be interpreted in an absolute sense, and they remain sensitive to the treatment of electrostatic interactions [72]. Nevertheless, it serves as a valuable qualitative tool in this study. The GQDs share similar poses, interacting most strongly with LYS317 and LYS321 of both protofilaments. These binding poses (Figure 5C) overlap with the location of the unidentified anionic density in the original cryo-electron microscopy structures [13], which Fitzpatrick *et al.* proposed may stabilize the interface. Indeed, we observe smaller RMSD values (by between 0.51 and 0.57 Å, Table 1) relative to the apo system, indicating that this binding mode somewhat stabilizes the structure across the studied timescale. Upon interacting with the functionalized GQDs, the distance between ARG349 and LYS375 decreases slightly in Protofilament 1 and by 1.04 to 4.66 Å in Protofilament 2 (Table 2). Moreover, the protofilaments twist relative to each other, which slightly narrows the large protofilament interface while broadening the small protofilament interface, resulting in an increased BSA. This is accompanied by a localized deformation of the backbone structure between LYS317 and LYS321 in Protofilament 2, which enables further electrostatic interactions with the GQD. Half functionalized *O*^-^, *S*^-^, and *COO*^-^ achieved BFE values ranging from -162 to -155, -157 to -112, and -125 to -114 kcalmol^-1^ respectively, showing that oxygen or sulfur edge groups significantly drive binding affinity to this interface and should be prioritized. It is unclear whether this fibril-stabilizing effect would persist at longer timescales; however, this binding mode may be an early intermediate in the disaggregation process [30]. This GQD-SF interaction should also be capable of accommodating larger GQDs than in the present study, which are more commonly employed experimentally. Given the high affinity and binding stability at this site, we predict that this is the dominant binding mode for anionic GQDs interacting with SFs, in agreement with the docking study.

At the PHF protofilament interface, the initial docked poses for pristine and half functionalized *O*^-^ (*S*^-^) GQDs were primarily edge-driven interactions, whereas with half functionalized *COO*^-^, the docked pose was slightly more buried in the protofilament interface’s crevice. One pristine replicate remained in the docked pose, one formed a purely edge interaction, and one assumed the binding pose that we previously identified [31], yielding BFE values of -11.60 ± 3.04, -16.33 ± 2.58, and -25.12 ± 3.72 kcalmol^-1^, respectively (Table S4). In all replicates of the half functionalized *O*^-^ (*S*^-^) GQD, the GQD left its initial docked pose and translated to the binding pose that the pristine GQD was previously identified to bind in (Figure 5D), yielding BFE values around -45 kcalmol^-1^ (-69 to -64 kcalmol^-1^), and interacting primarily with SER324, LEU325, GLY326, and ASN327. The half functionalized *S*^-^ GQD climbed the fibril axis and became more solvent-exposed. The half functionalized *COO*^-^ GQD’s left its somewhat buried docked pose in all replicates, sliding along one protofibril toward the termini in two replicates. These two replicates yielded variable BFE values between -78 and -66 kcalmol^-1^ and interacted mainly with LYS317 and LYS321. Remarkably, in the other replicate, this GQD left its binding site and became captured in the pinched interaction by Protofilament 2, achieving a BFE value of -79.15 ± 4.83 kcalmol^-1^, underscoring that the pinched pose is a robust binding conformation. Upon GQD binding at the PHF interface, the distance between ARG349 and LYS375 decreases (increases) in Protofilament 1 (2) relative to the apo system. Similar to the SF large protofilament interface systems, but to a lower magnitude, the functionalized GQDs decrease the protein’s backbone RMSD values (by between 0.17 and 0.45 Å, Table 1). Nevertheless, GQD functionalization does not appear to produce consistent or stable binding at this interface.

#### 3.3.5 General Trends in GQD-Tau Aggregate Simulations

The binding site preference for each GQD based on BFE values generally follows the trends in the docking study and favors the sites in order of their degree of positive charge. In the case of SF, the large protofilament interface is by far the most favorable site, followed by the C-shaped curve and the termini sites (which have similar BFE value ranges when comparing equal degrees of functionalization), and lastly the small protofilament interface (Figure 6). In PHFs, the docking study suggests a moderate preference for binding to the C-shaped curve, with BFE values indicating that this preference is slightly stronger, followed by the protofilament interface and the termini binding sites (Figure 6). The quarter functionalized GQDs are less affected by binding site electrostatics and almost always yield BFE values between -55 and -40 kcalmol^-1^. In both SFs and PHFs, half functionalized GQDs in the C-shaped curve consistently induce the pinched protofilament conformation, whereas this is rarely sampled with quarter functionalized GQDs. In this study, we find that anionic functionalization enables significantly more favorable GQD binding affinities, guiding the future development of tau-targeting GQDs as a potential therapeutic for Alzheimer’s disease.

**Figure 6:**
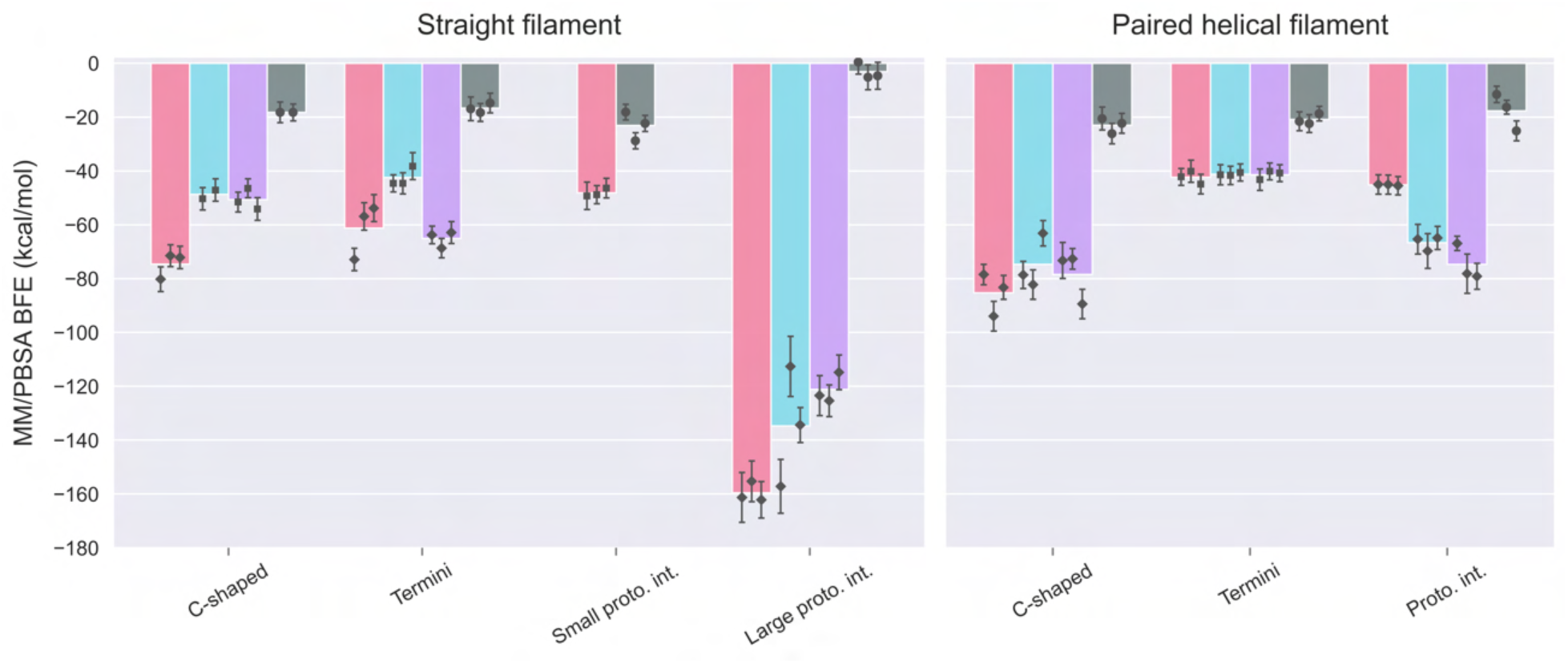
MM/PBSA BFE values across all systems. Bars represent the average BFE value among replicates, while individual markers show the mean value and standard deviation for each replicate. Bars within each binding site are plotted in the same order as in Table S4, where the leftmost bar is the first GQD listed in its binding site. The markers are also ordered by replicate, where replicate 1 is the leftmost. Marker shape indicates the degree of functionalization: diamonds (half), squares (quarter), and circles (none/pristine).

## 4 Conclusions

Growing evidence suggests that GQDs are capable of inhibiting the formation and disassembling TAs, with anionic GQDs being highly effective. In this study, we identified putative GQD binding sites in SF and PHF structures, then docked a library of edge-functionalized GQDs of various compositions and charge states to these sites. We found that the anionic GQDs yielded markedly more favorable docking scores than the other charge states. We selected a set of the most promising GQDs from each binding site and subjected them to MD simulations in triplicate, totaling 28 *µ*s of sampling. The anionic GQDs generally presented distinct interactions compared to the pristine (unfunctionalized) GQD, with a strong preference for the large protofilament interface in SFs, distorting the local protein structure, and a modest preference for the C-shaped curve in PHFs. In both TAs’ C-shaped curves, a unique protofilament conformation was discovered in which the GQD-bound protofilament underwent a constriction resembling a pinching motion and captured the GQD. This interaction is expected to destabilize TAs and can only be maintained in the presence of highly functionalized anionic GQDs. We also observed stable GQD surface interactions at the PHF6 region, which could prevent fibril elongation. Our work provides atomic insights into GQD-TA interactions and potential early mechanisms for GQD-mediated TA disassembly. Together, we propose that future experimental efforts focus on synthesis and preparation protocols that yield GQDs with a high degree of heteroatoms capable of possessing negative charges (*e.g.*, *COO*^-^, *O*^-^, and *S*^-^), with particular attention to sulfur, as it is less studied. This strategy should produce GQDs with enhanced potency, providing a more defined framework for the development of nanoparticle-based AD therapeutics.

## 5 Associated Content

The supporting information is available free of charge at website.com.

- Figures: Representative equilibration outputs, TA APBS electrostatic surfaces, distributions of docking scores by binding site, bound poses of GQDs selected for MD simulations, GQD scaffold, Protofilament 2 ARG349-LYS375 distance distributions, FESs and representative structures of apo TAs, linear regression between average MM/PBSA BFEs and docking scores.
- Tables: Docking scores for all GQDs, Z-scores and interactions for selected GQDs, MD TA summary statistics, MD TA differences and distribution overlaps between GQD-TA systems and apo systems, MM/PBSA BFEs and notable interactions.

## 6 Author Information

### 6.1 Corresponding Author

**Subha Kalyaanamoorthy** - Department of Chemistry, University of Waterloo, Waterloo, ON N2L 3G1, Canada;

### 6.2 Author

**Max Walton-Raaby** - Department of Chemistry, University of Waterloo, Waterloo, ON N2L 3G1, Canada;

Complete contact information is available at: website.com

## 8 Notes

The authors declare no competing financial interests.

## 9 Acknowledgments

This research was enabled in part by support provided by Compute Ontario (computeontario.ca) and the Digital Research Alliance of Canada (alliancecan.ca). M.W.-R thanks Paula Jofily for many helpful discussions regarding this project and assistance with constructing the graphical abstract.

