## Supplementary for "Computational Modeling of Functionalized Graphene Quantum Dots Binding to Tau Fibrils in Alzheimer’s Disease"

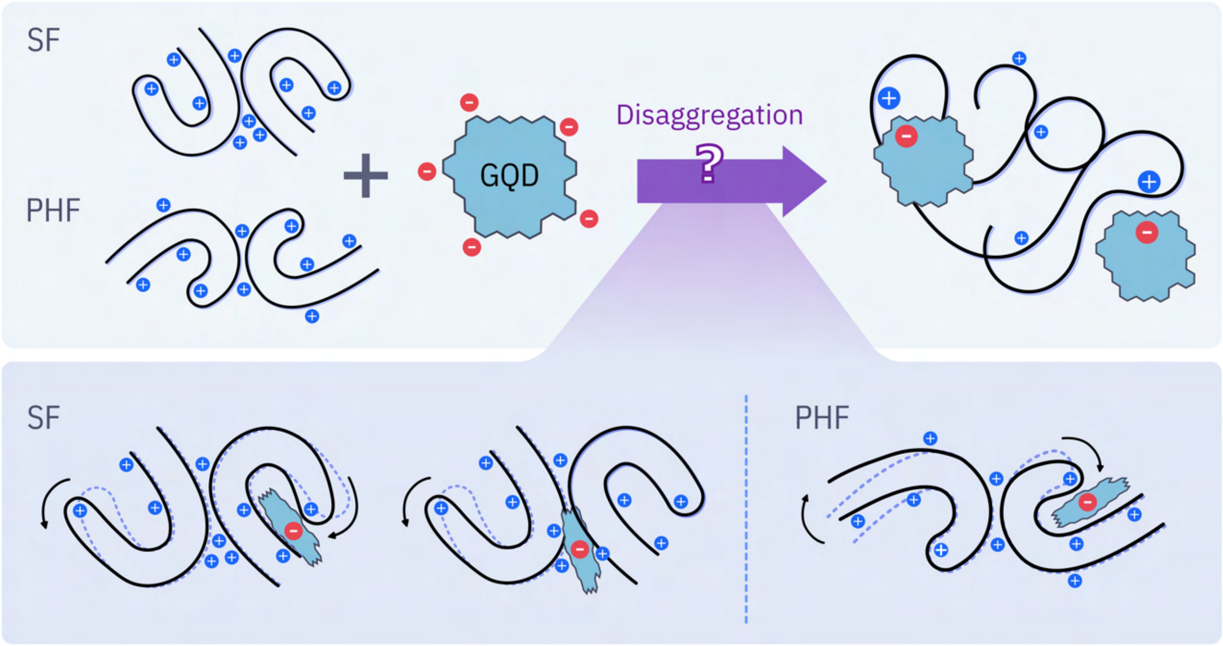


### Supplementary Information

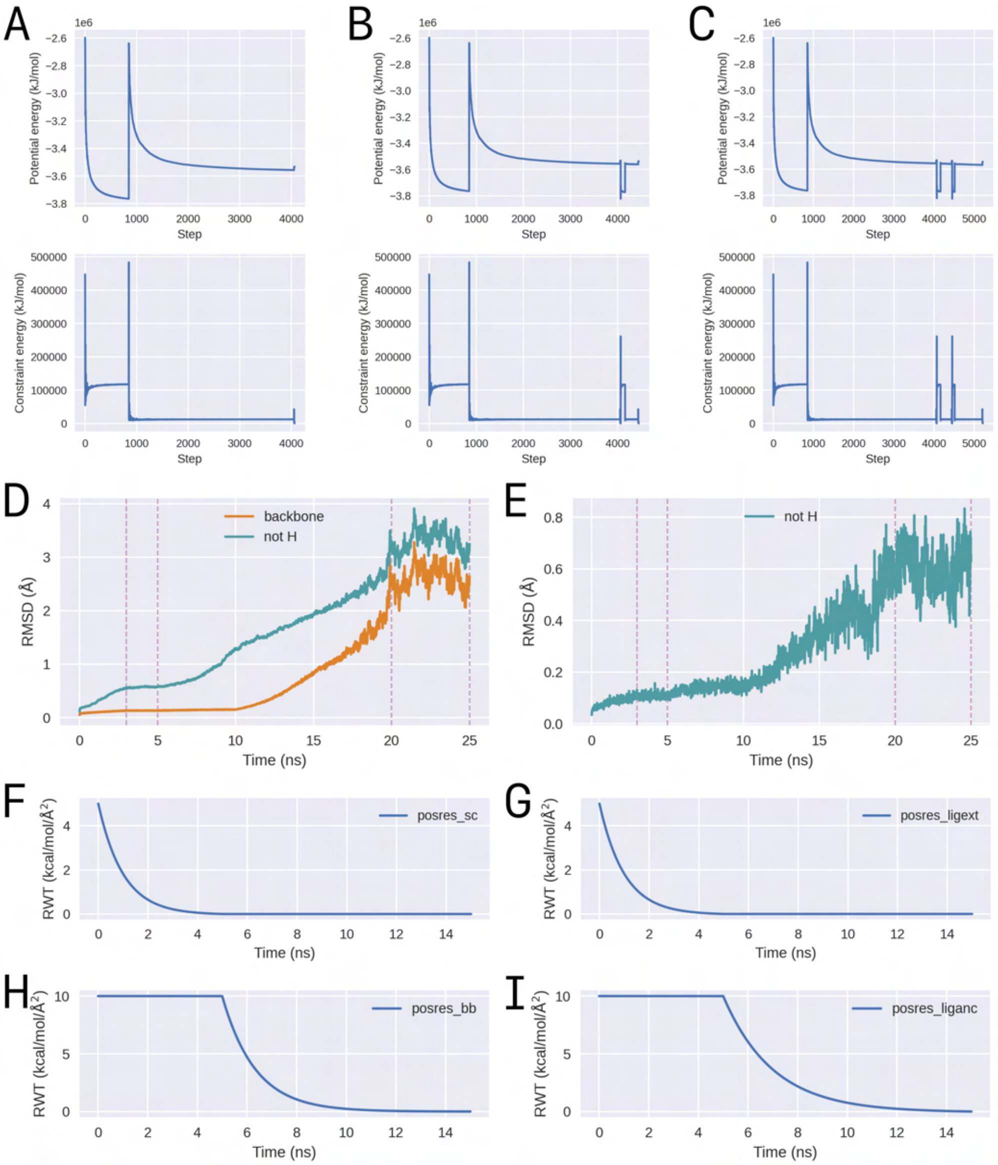


**Figure S1:** Summary of parameters and outputs from a representative 25 ns equilibration with MD-Protocol (Ref 54): 3 ns heating, 2 ns NVT, 15 ns NPT with restraints (NPTr), and 5 ns NPT with no restraints. Minimization stages 1 (A), 2 (B), and 3 (C); Protein backbone and heavy atom root-mean square deviation (RMSD) where restraint weights are adjusted between 5 and 20 ns (D); Ligand heavy atom RMSD where restraint weights are adjusted between 5 and 20 ns (E); Protein side chain restraint weight in NPTr (F); Ligand extension restraint weight in NPTr (G); Protein backbone restraint weight in NPTr (H); Ligand anchor restraint weight in NPTr (I).

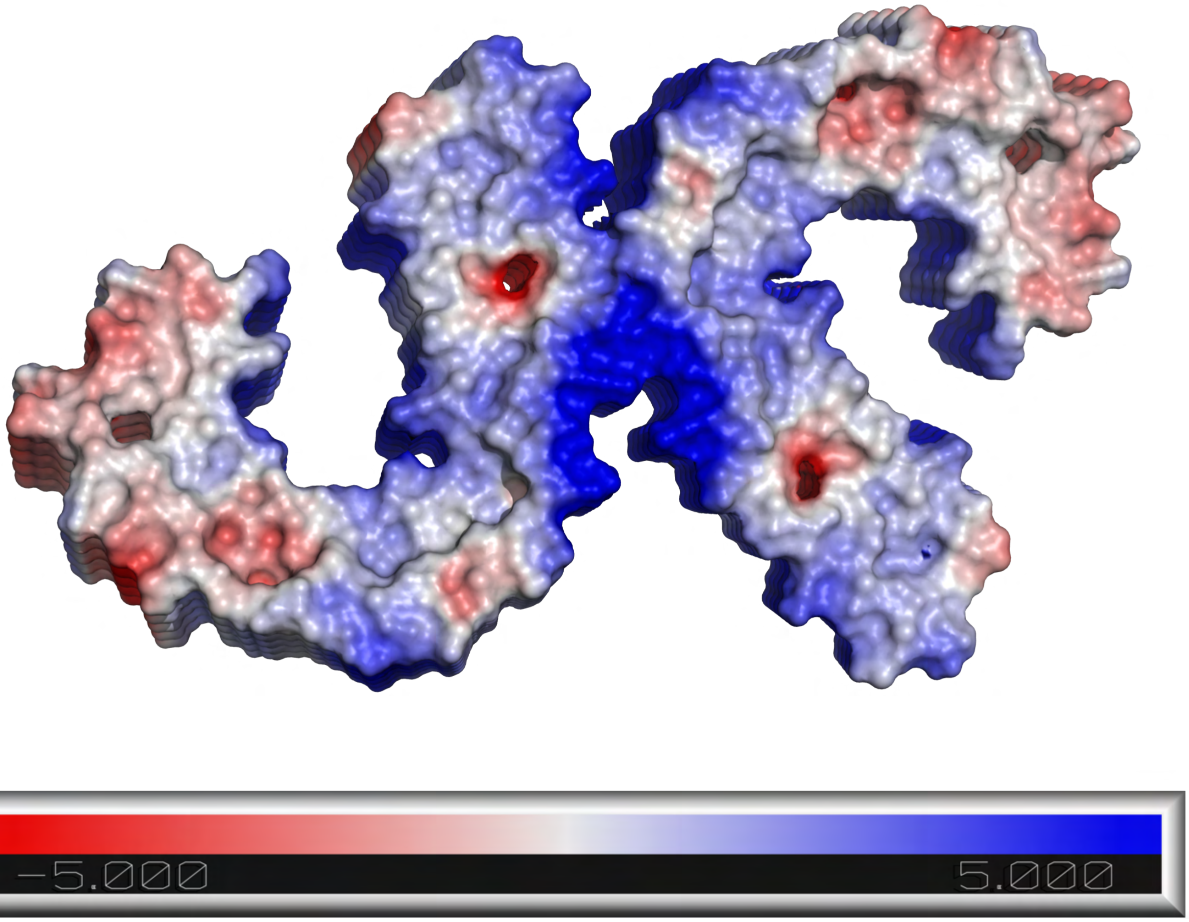


**Figure S2:** Straight filament (SF) electrostatic surface calculated with the Adaptive Poisson-Boltzmann Solver method.

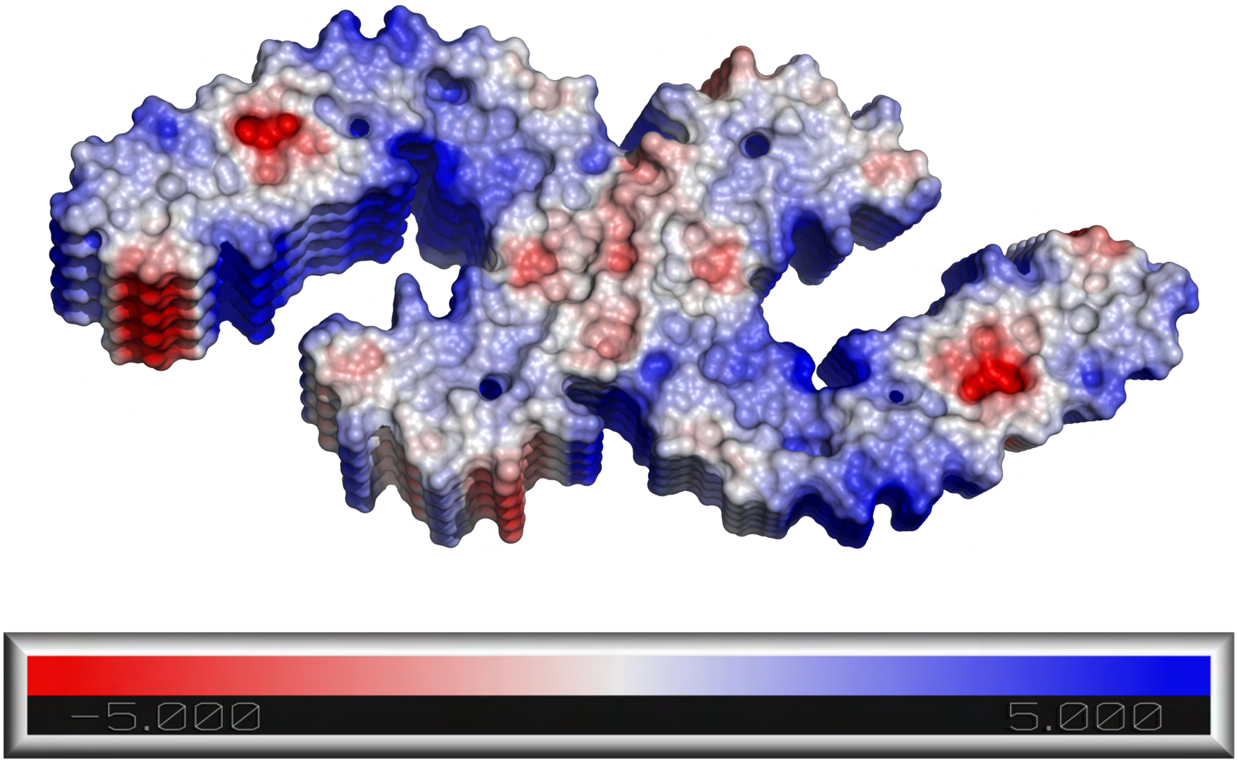


**Figure S3:** Paired helical filament (PHF) electrostatic surface calculated with the Adaptive Poisson-Boltzmann Solver method.

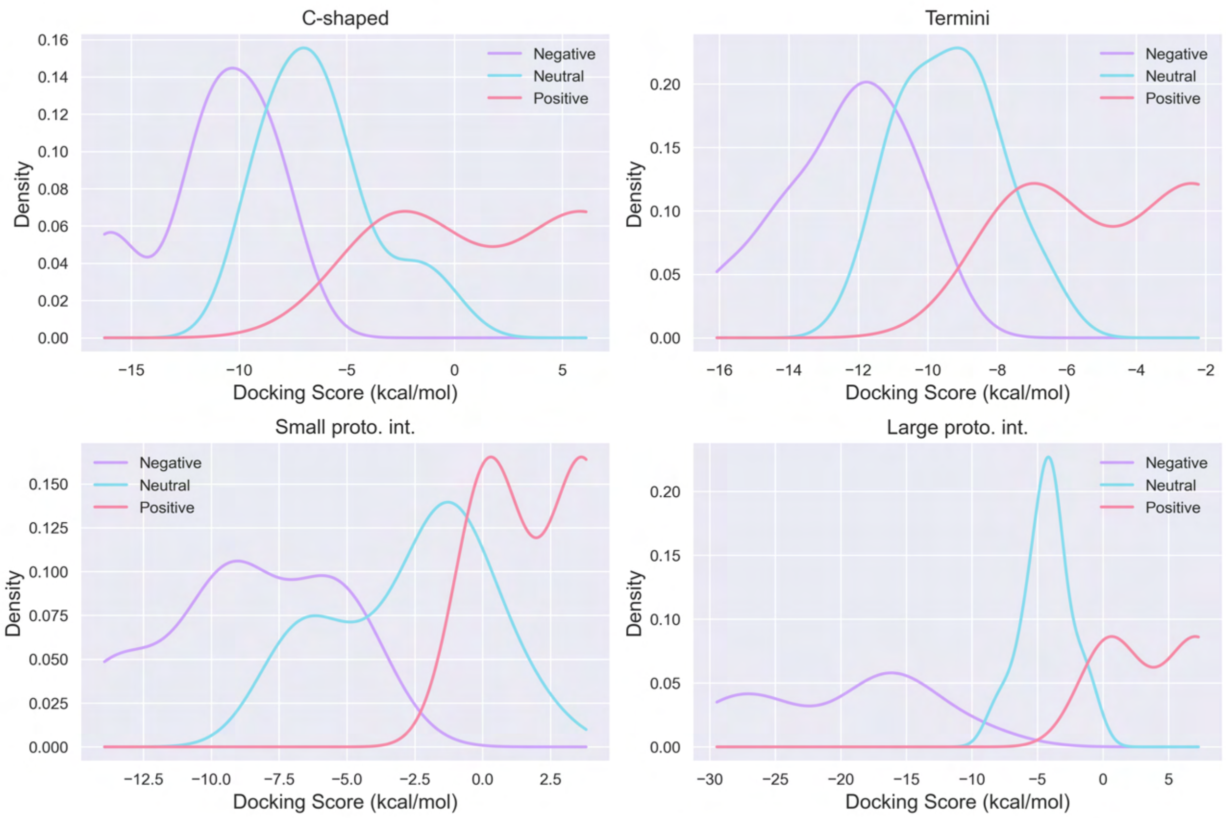


**Figure S4:** Kernel density estimate (KDE) of SF docking scores in identified binding sites.

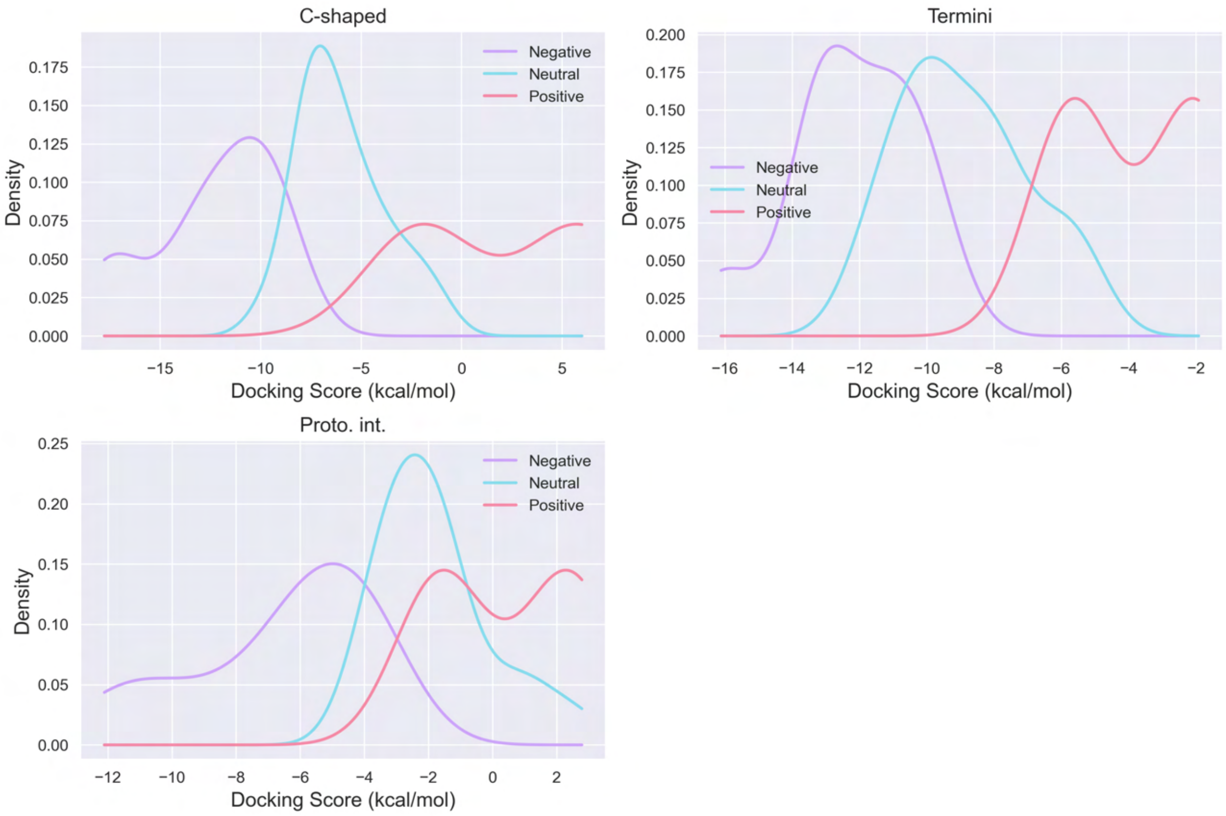


**Figure S5:** KDE of PHF docking scores in identified binding sites.

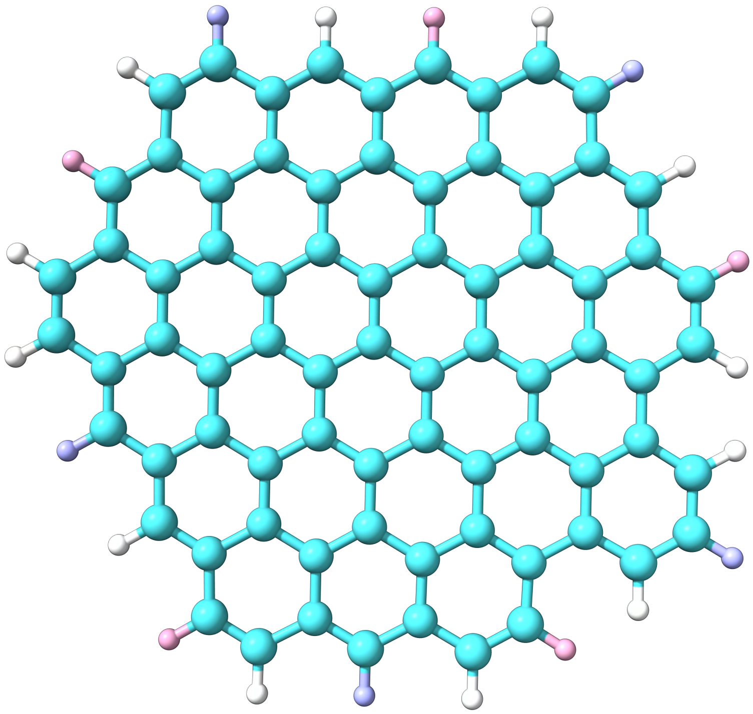

**Figure S6:** Structure of the studied graphene quantum dot (GQD). Quarter functionalization GQDs are modified at the pink groups, while half-functionalization GQDs are modified at both pink and purple groups. The pristine GQD has hydrogen atoms at all edge groups.

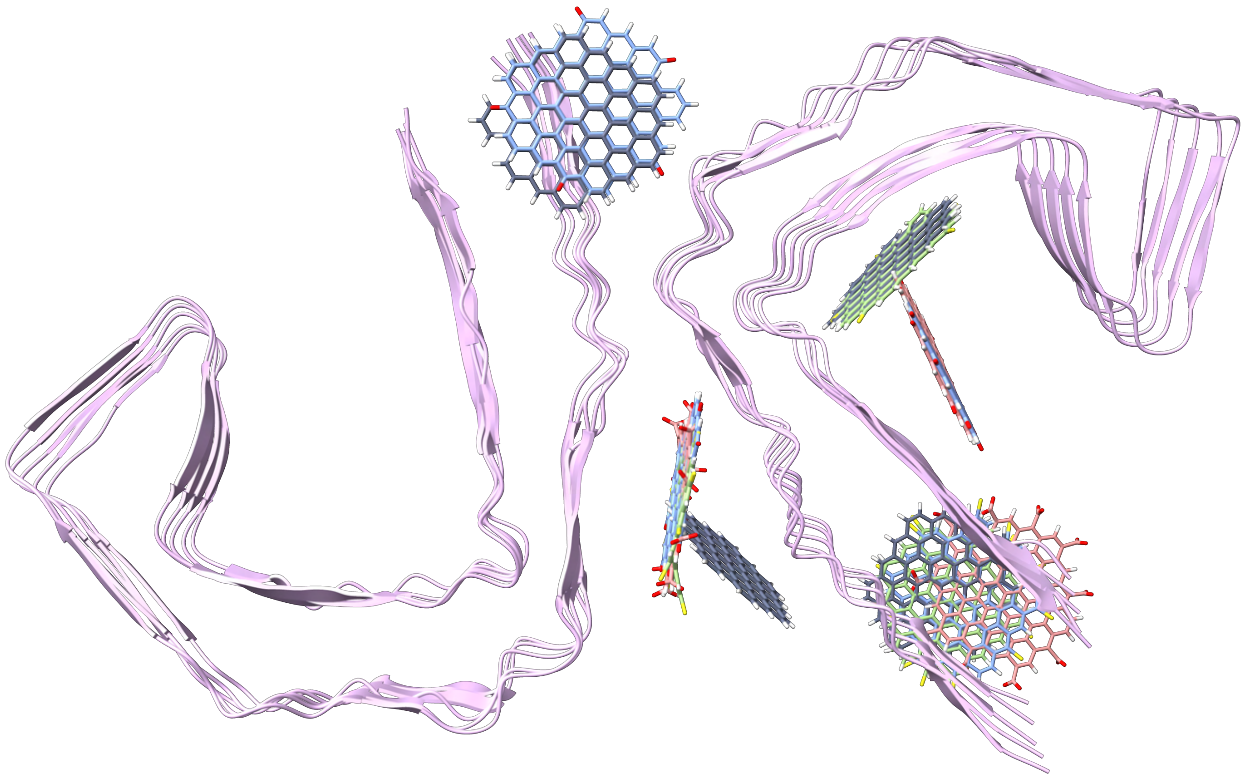

**Figure S7:** Selected GQDs for molecular dynamics (MD) simulations with SF. Functionalized GQDs are colored in ascending order of docking scores in each binding site: blue (best / most negative docking score), green (second best), and red (third best), while the pristine GQD is shown in gray. The specific GQDs are listed in Table S3.

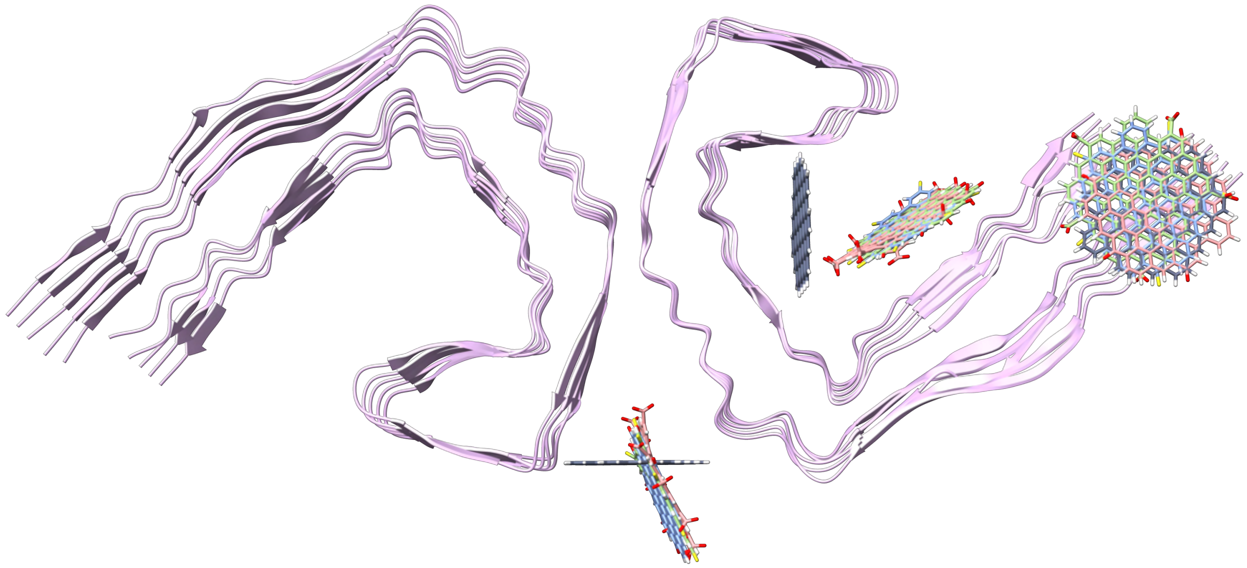

**Figure S8:** Selected GQDs for MD simulations with PHF. Functionalized GQDs are colored in ascending order of docking scores in each binding site: blue (best / most negative docking score), green (second best), and red (third best), while the pristine GQD is shown in gray. The specific GQDs are listed in Table S3.

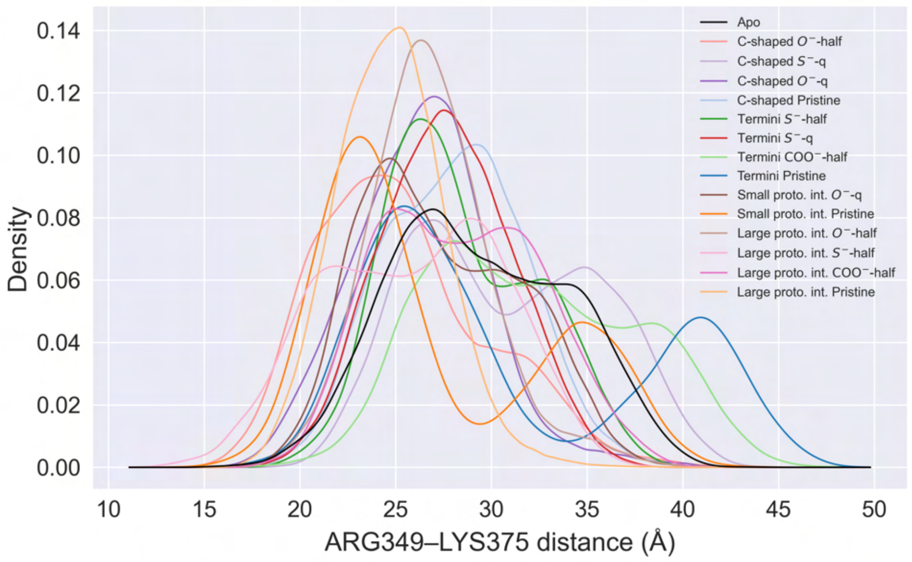


**Figure S9:** KDE of ARG349–LYS375 distance distributions for SF Protofilament 2.

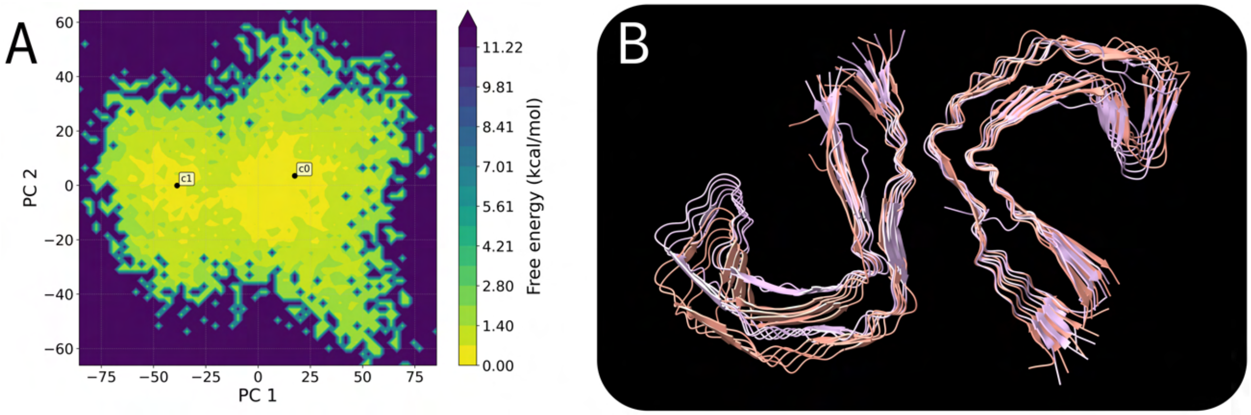


**Figure S10:** Free energy landscapes with respect to principle components (PC) 1 and 2 for apo SF (A), and the most representative structure from cluster 0 (c0, pink) and cluster 1 (c1, salmon) (B).

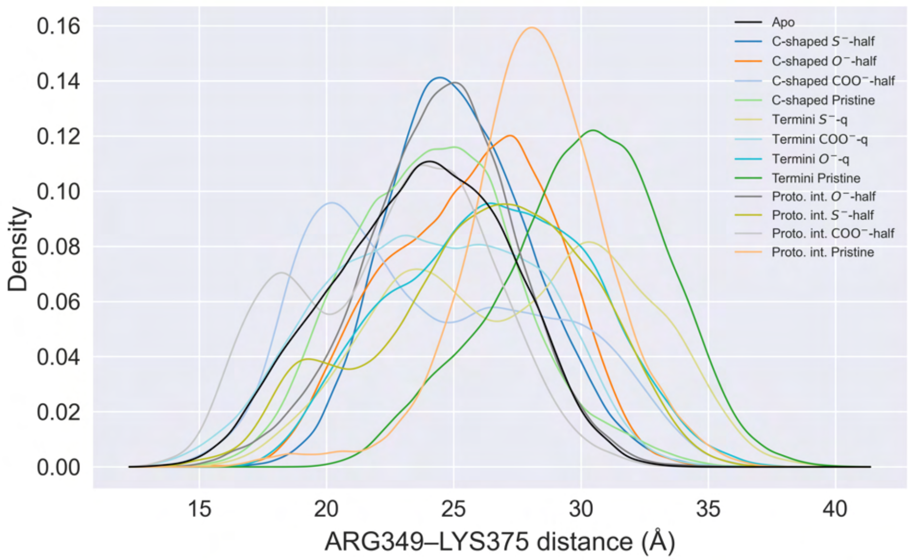


**Figure S11:** KDE of ARG349–LYS375 distance distributions for PHF Protofilament 2.

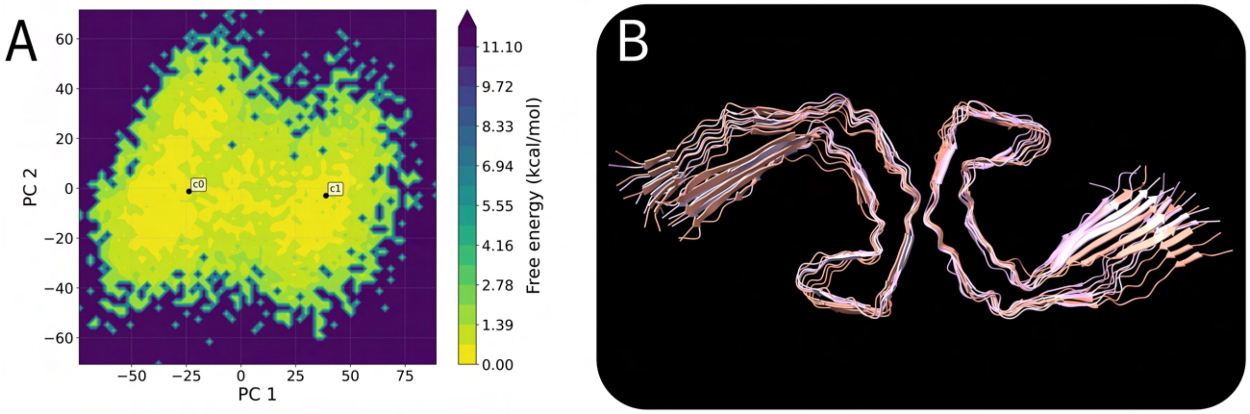


**Figure S12:** Free energy landscapes with respect to PC1 and PC2 for apo PHF (A), and the most representative structure from cluster 0 (c0, pink) and cluster 1 (c1, salmon) (B).

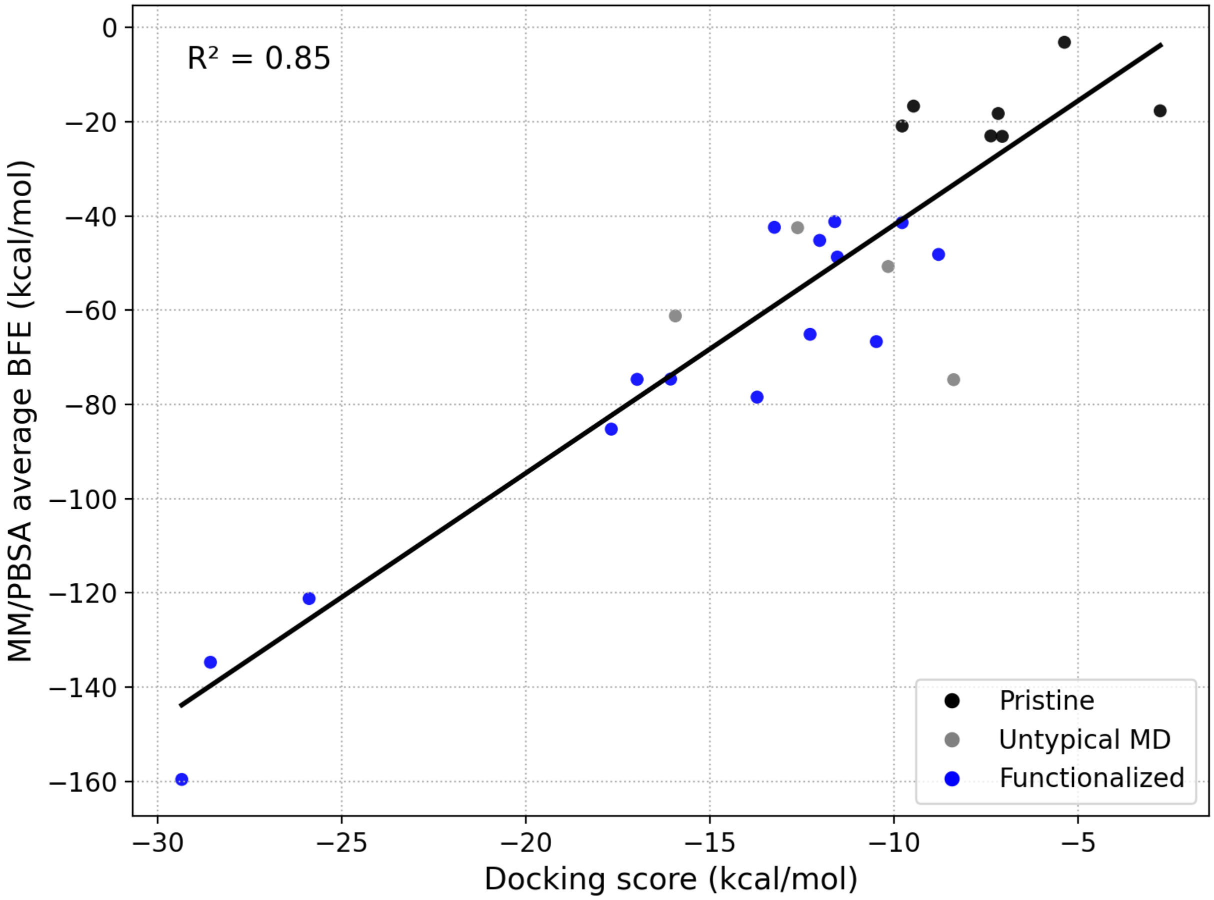


**Figure S13:** Linear regression between the average molecular mechanics Poisson-Boltzmann surface area (MM/PBSA) binding free energy (BFE) value across replicates and the docking score of the starting pose. Black points represent pristine systems, blue points represent functionalized systems, and grey points represent systems that had large variation in replicates (*e.g.*, due to the pinched protofilament conformation).

Table S1: Docking scores of GQDs binding to straight filament.

| Binding site | GQD | Docking scores ($kcal\,mol^{-1}$) |
|---|---|---|
| **C-shaped** | $O^-$-half | −16.14 |
| | $S^-$-half | −15.92 |
| | $S^-$-q | −11.53 |
| | $COO^-$-half | −11.44 |
| | $O^-$-q | −10.33 |
| | $CH_3$-half | −9.72 |
| | NO-q | −9.63 |
| | $COO^-$-q | −8.71 |
| | $OH-COO^-$-q | −8.70 |
| | NO-half | −8.59 |
| | $CH_3$-q | −8.47 |
| | SH-q | −8.44 |
| | $OH-COO^-$-half | −7.80 |
| | F-half | −7.57 |
| | Pristine | −7.17 |
| | F-q | −7.15 |
| | SH-half | −6.84 |
| | $OCH_3$-half | −6.21 |
| | $OCH_3$-q | −6.19 |
| | OH-q | −5.86 |
| | OH-COOH-q | −5.73 |
| | $NH_2$-q | −5.04 |
| | OH-half | −5.00 |
| | COOH-q | −4.02 |
| | $NH_3^+$-q | −2.47 |
| | $NH_2$-half | −1.90 |
| | OH-COOH-half | −1.75 |
| | COOH-half | −0.22 |
| | $NH_3^+$-half | 6.01 |
| **Termini** | $S^-$-half | −15.97 |
| | $O^-$-half | −14.18 |
| | $S^-$-q | −12.62 |

| | | |
|---|---|---|
| | $COO^-$-half | −12.33 |
| | $CH_3$-half | −11.96 |
| | $COO^-$-q | −11.62 |
| | $CH_3$-q | −10.94 |
| | NO-q | −10.91 |
| | $O^-$-q | −10.89 |
| | $OH-COO^-$-half | −10.57 |
| | SH-half | −10.48 |
| | SH-q | −10.44 |
| | NO-half | −9.92 |
| | $OH-COO^-$-q | −9.79 |
| | F-half | −9.73 |
| | F-q | −9.55 |
| | Pristine | −9.47 |
| | OH-q | −9.36 |
| | COOH-q | −8.77 |
| | $OCH_3$-q | −8.70 |
| | $NH_2$-q | −8.65 |
| | OH-half | −8.41 |
| | OH-COOH-q | −8.28 |
| | $OCH_3$-half | −8.26 |
| | OH-COOH-half | −7.45 |
| | $NH_2$-half | −7.14 |
| | $NH_3^+$-q | −7.04 |
| | COOH-half | −6.41 |
| | $NH_3^+$-half | −2.31 |
| **Small proto. int.** | $S^-$-half | −13.83 |
| | $O^-$-half | −13.12 |
| | $COO^-$-half | −9.97 |
| | $S^-$-q | −9.43 |
| | $O^-$-q | −9.09 |
| | $CH_3$-q | −7.46 |
| | Pristine | −7.26 |
| | F-q | −6.65 |
| | NO-q | −6.43 |
| | $COO^-$-q | −6.32 |

| | | |
|---|---|---|
| | F-half | −6.10 |
| | $OH-COO^-$-half | −5.34 |
| | $CH_3$-half | −4.85 |
| | $OH-COO^-$-q | −4.32 |
| | OH-q | −4.18 |
| | $OCH_3$-q | −3.43 |
| | NO-half | −2.34 |
| | OH-half | −2.31 |
| | SH-q | −1.75 |
| | $NH_2$-q | −1.44 |
| | SH-half | −1.05 |
| | OH-COOH-q | −0.93 |
| | $OCH_3$-half | −0.83 |
| | COOH-q | −0.29 |
| | $NH_3^+$-q | 0.23 |
| | $NH_2$-half | 0.58 |
| | OH-COOH-half | 0.73 |
| | COOH-half | 2.23 |
| | $NH_3^+$-half | 3.71 |
| **Large proto. int.** | $O^-$-half | −29.35 |
| | $S^-$-half | −28.56 |
| | $COO^-$-half | −26.00 |
| | $COO^-$-q | −17.46 |
| | $S^-$-q | −17.38 |
| | $O^-$-q | −16.63 |
| | $OH-COO^-$-half | −14.36 |
| | $OH-COO^-$-q | −9.56 |
| | NO-half | −8.30 |
| | NO-q | −7.40 |
| | $CH_3$-half | −6.29 |
| | $CH_3$-q | −5.99 |
| | Pristine | −5.37 |
| | F-q | −5.21 |
| | F-half | −5.15 |
| | $OCH_3$-q | −4.69 |
| | SH-q | −4.47 |

| | |
|---|---|
| SH-half | −3.98 |
| $OCH_3$-half | −3.96 |
| OH-q | −3.84 |
| OH-COOH-q | −3.74 |
| COOH-q | −3.61 |
| $NH_2$-q | −3.60 |
| OH-half | −2.40 |
| $NH_2$-half | −1.82 |
| OH-COOH-half | −1.45 |
| COOH-half | −0.64 |
| $NH_3^+$-q | 0.53 |
| $NH_3^+$-half | 7.19 |

Table S2: Docking scores of GQDs binding to paired helical filament.

| Binding site | GQD | Docking scores (kcal mol$^{-1}$) |
|---|---|---|
| **C-shaped** | $S^-$-half | −17.67 |
| | $O^-$-half | −16.98 |
| | $COO^-$-half | −13.80 |
| | $S^-$-q | −11.93 |
| | $O^-$-q | −11.26 |
| | $COO^-$-q | −10.29 |
| | NO-q | −9.65 |
| | $OH-COO^-$-half | −9.15 |
| | $OH-COO^-$-q | −8.71 |
| | $CH_3$-q | −8.21 |
| | $CH_3$-half | −7.81 |
| | F-half | −7.77 |
| | F-q | −7.61 |
| | NO-half | −7.41 |
| | Pristine | −7.36 |
| | SH-q | −7.00 |
| | OH-q | −6.70 |
| | OH-COOH-q | −6.51 |
| | $OCH_3$-q | −5.88 |
| | OH-half | −5.87 |
| | $NH_2$-q | −5.68 |
| | SH-half | −5.43 |
| | $OCH_3$-half | −4.61 |
| | COOH-q | −4.00 |
| | $NH_2$-half | −2.83 |
| | OH-COOH-half | −2.17 |
| | $NH_3^+$-q | −2.02 |
| | COOH-half | −1.21 |
| | $NH_3^+$-half | 5.89 |
| **Termini** | $S^-$-half | −16.01 |
| | $O^-$-half | −13.39 |
| | $S^-$-q | −13.26 |

| | | |
|---|---|---|
| | $COO^-$-half | −12.59 |
| | $CH_3$-half | −11.74 |
| | $COO^-$-q | −11.70 |
| | NO-q | −11.56 |
| | $CH_3$-q | −11.05 |
| | NO-half | −10.98 |
| | $O^-$-q | −10.57 |
| | $OH-COO^-$-half | −10.37 |
| | F-half | −10.18 |
| | SH-q | −9.98 |
| | SH-half | −9.86 |
| | Pristine | −9.84 |
| | F-q | −9.81 |
| | $OH-COO^-$-q | −9.57 |
| | $OCH_3$-q | −8.70 |
| | OH-q | −8.24 |
| | OH-COOH-q | −8.05 |
| | $OCH_3$-half | −7.90 |
| | $NH_2$-q | −7.89 |
| | COOH-q | −7.88 |
| | OH-half | −6.68 |
| | OH-COOH-half | −5.78 |
| | $NH_3^+$-q | −5.67 |
| | $NH_2$-half | −5.66 |
| | COOH-half | −5.43 |
| | $NH_3^+$-half | −2.02 |
| **Proto. int.** | $O^-$-half | −12.01 |
| | $S^-$-half | −10.49 |
| | $COO^-$-half | −8.50 |
| | $O^-$-q | −6.32 |
| | $S^-$-q | −5.54 |
| | $OH-COO^-$-q | −4.67 |
| | $COO^-$-q | −4.56 |
| | NO-q | −4.12 |
| | $CH_3$-half | −3.80 |
| | NO-half | −3.77 |

| | |
|---|---|
| $OH-COO^{-}$-half | −3.52 |
| $CH_3$-q | −3.24 |
| F-half | −2.90 |
| F-q | −2.84 |
| Pristine | −2.76 |
| SH-q | −2.60 |
| OH-q | −2.46 |
| $NH_2$-q | −1.88 |
| OH-COOH-q | −1.86 |
| $OCH_3$-q | −1.83 |
| $NH_3^{+}$-q | −1.60 |
| SH-half | −1.46 |
| OH-half | −1.38 |
| $OCH_3$-half | −1.05 |
| COOH-q | 0.12 |
| $NH_2$-half | 0.84 |
| OH-COOH-half | 1.43 |
| $NH_3^{+}$-half | 2.37 |
| COOH-half | 2.69 |

Table S3: Docking scores, Z-scores, and interactions of selected GQDs, including chain letter.

| Aggregate | Binding site | GQD | Docking scores ($kcal\,mol^{-1}$) | Local Z-score | Average Z-score | Interactions |
|---|---|---|---|---|---|---|
| **Straight filament** | **C-shaped** | $O^-$-half | −16.14 | −2.06 | −2.05 | C:LYS369, C:ILE371, C:THR373, E:LYS369, E:ILE371, G:LYS369, G:ILE371, I:LYS369 |
| | | $S^-$-q | −11.53 | −1.01 | −1.03 | A:ILE360, C:LYS353, C:ILE360, C:LYS369, E:ASP358, E:ILE360, E:LYS369, G:ILE360, G:LYS369 |
| | | $O^-$-q | −10.33 | −0.69 | −0.75 | C:LYS369, C:ILE371, C:THR373, E:LYS369, E:ILE371, G:LYS369, I:LYS369 |
| | | Pristine | −7.17 | | | C:ASP358, C:ILE360, C:LYS369, E:ILE360, G:ILE360 |
| | **Termini** | $S^-$-half | −15.97 | −2.60 | −2.24 | A:ILE308, A:TYR310, A:LEU376, A:PHE378 |
| | | $S^-$-q | −12.62 | −1.26 | −1.03 | A:ILE308, A:TYR310, A:LYS311, A:LEU376, A:PHE378 |
| | | $COO^-$-half | −12.33 | −1.13 | −1.28 | A:VAL306, A:ILE308, A:TYR310, A:LYS375, A:LEU376, A:PHE378 |
| | | Pristine | −9.47 | | | A:TYR310, A:LYS311, A:LEU376, A:PHE378 |

| | | | | | | |
|---|---|---|---|---|---|---|
| | **Small proto. int.** | $O^-$-q | −9.09 | −0.41 | −0.75 | J:VAL306, J:GLN307, J:VAL309, J:LYS311 |
| | | Pristine | −7.26 | | | J:VAL306, J:GLN307, J:VAL309, J:TYR310 |
| | **Large proto. int.** | $O^-$-half | −29.35 | −2.82 | −2.05 | C:LYS317, C:THR319, C:LYS321, D:LYS317, D:LYS321, E:LYS317, E:THR319, E:LYS321, F:LYS317, F:LYS321, G:LYS317, G:THR319, H:LYS317, H:LYS321, I:LYS317 |
| | | $S^-$-half | −28.56 | −2.73 | −2.24 | C:LYS317, E:LYS317, E:THR319, F:LYS317, F:LYS321, G:LYS317, G:THR319, J:LYS321 |
| | | $COO^-$-half | −26.00 | −2.41 | −1.28 | A:LYS317, A:LYS321, B:LYS317, C:LYS317, C:LYS321, D:LYS317, D:LYS321, E:LYS317, E:THR319, E:LYS321, F:LYS317, F:LYS321, G:LYS317, H:LYS317, H:LYS321, I:LYS317, J:LYS321 |
| | | Pristine | −5.37 | | | A:LEU315, C:LEU315, C:LYS317, E:LEU315, G:LEU315 |
| **Paired helical filament** | **C-shaped** | $S^-$-half | −17.67 | −2.25 | −2.26 | C:ILE371, E:ILE371, E:THR373, G:LYS369, G:ILE371, G:THR373 |
| | | $O^-$-half | −16.98 | −2.10 | −2.06 | C:LYS369, C:ILE371, E:LYS369, E:ILE371, E:THR373, G:LYS369, G:ILE371 |

| | | | | | |
|---|---|---|---|---|---|
| | $COO^-$-half | −13.80 | −1.39 | −1.34 | A:LYS369, C:LYS369, C:ILE371, E:LYS369, E:ILE371, E:THR373, G:LYS369, G:ILE371, G:THR373, G:LYS375, I:LYS369 |
| | Pristine | −7.36 | | | A:ILE360, C:LYS353, C:ILE360, E:LYS353, E:ILE360, G:LYS353, G:ILE360 |
| **Termini** | $S^-$-q | −13.26 | −1.23 | −1.01 | A:VAL306, A:ILE308, A:VAL309, A:TYR310, A:LEU376, A:PHE378 |
| | $COO^-$-q | −11.70 | −0.65 | −0.60 | A:VAL306, A:GLN307, A:ILE308, A:TYR310, A:LYS311, A:LYS375, A:LEU376 |
| | $O^-$-q | −10.57 | −0.27 | −0.73 | A:VAL306, A:GLN307, A:ILE308, A:VAL309, A:TYR310, A:LEU376 |
| | Pristine | −9.84 | | | A:VAL306, A:ILE308, A:VAL309, A:TYR310, A:LEU376 |
| **Proto. int.** | $O^-$-half | −12.01 | −2.82 | −2.06 | D:LYS340, F:LYS340, H:LYS340, J:LYS340 |
| | $S^-$-half | −10.49 | −2.35 | −2.26 | J:LYS340 |
| | $COO^-$-half | −8.50 | −1.71 | −1.34 | B:LYS340, C:LYS331, D:LYS340, F:LYS340, H:LYS340, J:LYS340 |
| | Pristine | −2.76 | | | F:LYS340, F:GLU342, H:LYS340, H:GLU342, J:LYS340 |

Table S4: MM/PBSA BFE values for each replicate, with the top three electrostatic and van der Waals interactions listed.

| Aggregate | Binding site | GQD | Rep. # | ΔG ($\mathrm{kcal\,mol^{-1}}$) | Electrostatic interactions | vdW interactions |
|---|---|---|---|---|---|---|
| **Straight filament** | **C-shaped** | $O^-$-half | 1 | $-80.22 \pm 4.61$ | ARG349, LYS375, LYS369 | GLN351, ILE371, THR373 |
| | | | 2 | $-71.49 \pm 4.06$ | ARG349, LYS369, LYS375 | GLN351, ILE371, HIS362 |
| | | | 3 | $-72.15 \pm 4.12$ | ARG349, LYS375, LYS347 | GLN351, THR373, ILE371 |
| | | $S^-$-q | 1 | Outlier | | |
| | | | 2 | $-50.34 \pm 4.16$ | LYS353, LYS369 | HIS362, ILE360, HIS330 |
| | | | 3 | $-47.11 \pm 4.16$ | LYS369, LYS353, LYS370 | HIS362, PRO364, ILE360 |
| | | $O^-$-q | 1 | $-51.56 \pm 3.71$ | LYS369, LYS375, LYS370 | ILE371, THR373, HIS362 |
| | | | 2 | $-46.48 \pm 3.51$ | LYS353, ARG349, LYS375 | GLN351, SER352, VAL350 |
| | | | 3 | $-54.16 \pm 4.21$ | LYS369, LYS375, LYS370 | ILE371, THR373, HIS362 |
| | | Pristine | 1 | $-18.25 \pm 3.81$ | ASP358, LYS369, LYS353 | HIS362, ILE360, PRO364 |
| | | | 2 | Outlier | | |

| | | | | | |
|---|---|---|---|---|---|
| | | 3 | −18.27 ± 3.13 | LYS369, ASP358, LYS331 | HIS362, ILE360, PRO364 |
| **Termini** | $S^-$-half | 1 | −72.92 ± 4.14 | ARG349, LYS375, LYS353 | GLN351, THR377, VAL350 |
| | | 2 | −56.89 ± 5.10 | NVAL306, LYS311, LYS375 | GLN307, THR377, VAL309 |
| | | 3 | −53.84 ± 4.97 | LYS375, LYS311, NVAL306 | GLN307, THR373, HIS374 |
| | $S^-$-q | 1 | −44.58 ± 3.18 | NVAL306, LYS311, LYS375 | GLN307, VAL309, LEU376 |
| | | 2 | −44.65 ± 3.92 | NVAL306, LYS311, LYS375 | GLN307, VAL309, LEU376 |
| | | 3 | −38.25 ± 5.00 | LYS375, LYS311, NVAL306 | THR373, TYR310, LEU376 |
| | $COO^-$-half | 1 | −63.78 ± 3.27 | NVAL306, LYS375, LYS311 | GLN307, VAL309, THR377 |
| | | 2 | −68.69 ± 3.63 | NVAL306, LYS311, LYS375 | GLN307, VAL309, LEU376 |
| | | 3 | −62.89 ± 4.03 | LYS375, ARG349, NVAL306 | THR373, HIS374, THR377 |
| | Pristine | 1 | −16.96 ± 4.37 | CPHE378, NVAL306, LYS375 | LEU376, TYR310, CPHE378 |
| | | 2 | −18.32 ± 3.30 | CPHE378, NVAL306, LYS375 | LEU376, CPHE378, ILE308 |

| | | | | | |
|---|---|---|---|---|---|
| | | 3 | −14.79 ± 3.61 | CPHE378, NVAL306, LYS375 | CPHE378, ILE308, LEU376 |
| **Small proto. int.** | $O^-$-q | 1 | −49.26 ± 5.12 | NVAL306, LYS311, LYS331 | GLN307, HIS329, SER324 |
| | | 2 | −48.86 ± 3.36 | LYS311, NVAL306, LYS370 | PRO312, GLN307, TYR310 |
| | | 3 | −46.40 ± 3.62 | LYS311, NVAL306, LYS370 | PRO312, TYR310, ILE308 |
| | Pristine | 1 | −18.15 ± 2.93 | LYS311, CPHE378, ASP314 | LYS311, PRO312, TYR310 |
| | | 2 | −28.81 ± 3.00 | LYS375, CPHE378, LYS311 | TYR310, LYS375, LEU376 |
| | | 3 | −22.36 ± 3.02 | LYS311, CPHE378, LYS375 | TYR310, ILE308, LYS311 |
| **Large proto. int.** | $O^-$-half | 1 | −161.31 ± 9.26 | LYS317, LYS321 | THR319, ASN368 |
| | | 2 | −155.28 ± 7.58 | LYS317, LYS321 | THR319, ASN368, SER316 |
| | | 3 | −162.18 ± 6.81 | LYS317, LYS321 | THR319, ASN368, SER316 |
| | $S^-$-half | 1 | −157.18 ± 10.00 | LYS321, LYS317 | THR319, ASN368, SER316 |
| | | 2 | −112.64 ± 11.14 | LYS321, LYS317 | THR319, SER316, ASN368 |

| | | | | | |
|---|---|---|---|---|---|
| | | 3 | $-134.36 \pm 6.50$ | LYS317, LYS321 | THR319, ASN368, SER316 |
| | $COO^{-}$-half | 1 | $-123.43 \pm 7.43$ | LYS317, LYS321 | THR319, ASN368, SER316 |
| | | 2 | $-125.37 \pm 5.91$ | LYS317, LYS321 | THR319, ASN368, SER316 |
| | | 3 | $-114.82 \pm 6.44$ | LYS317, LYS321 | THR319, ASN368 |
| | Pristine | 1 | $0.44 \pm 4.50$ | LYS321, ASP314, GLU372 | LYS321, LEU315, VAL313 |
| | | 2 | $-5.20 \pm 4.63$ | LYS321, ASP314, GLU372 | LYS321, LEU315, VAL313 |
| | | 3 | $-4.66 \pm 5.03$ | LYS321, ASP314, GLU372 | LYS321, LEU315, VAL313 |
| **Paired helical filament** | **C-shaped** | | | | |
| | $S^{-}$-half | 1 | $-78.47 \pm 3.79$ | ARG349, LYS375, LYS369 | GLN351, HIS362, ILE371 |
| | | 2 | $-93.98 \pm 5.48$ | LYS369, ARG349, LYS375 | ILE371, GLN351, THR373 |
| | | 3 | $-83.29 \pm 4.43$ | LYS369, LYS375, LYS370 | ILE371, THR373, GLY367 |
| | $O^{-}$-half | 1 | $-78.63 \pm 5.08$ | ARG349, LYS369, LYS375 | ILE371, PRO364, ASN368 |
| | | 2 | $-82.24 \pm 5.50$ | ARG349, LYS375, LYS369 | GLN351, THR373, ILE371 |

| | | | | | |
|---|---|---|---|---|---|
| | | 3 | −63.17 ± 4.75 | LYS369, ARG349, LYS375 | GLN351, GLY367, ILE371 |
| | $COO^-$-half | 1 | −73.29 ± 6.66 | LYS369, LYS375, LYS370 | THR373, ILE371, GLY367 |
| | | 2 | −72.64 ± 3.83 | LYS369, LYS375, ARG349 | ILE371, THR373, HIS362 |
| | | 3 | −89.43 ± 5.48 | LYS369, ARG349, LYS375 | ILE371, GLN351, GLY367 |
| | Pristine | 1 | −20.56 ± 4.27 | LYS353, LYS369, ARG349 | LYS353, ILE360, GLN351 |
| | | 2 | −26.16 ± 3.75 | LYS353, LYS369, ARG349 | LYS353, ILE360, GLN351 |
| | | 3 | −22.34 ± 3.72 | LYS353, LYS369, ASP345 | LYS353, ILE360, GLN351 |
| **Termini** | $S^-$-q | 1 | −42.21 ± 3.11 | LYS311, NVAL306, LYS375 | TYR310, PRO312, ILE308 |
| | | 2 | −40.11 ± 4.11 | LYS311, NVAL306, LYS375 | ILE308, TYR310, LEU376 |
| | | 3 | −44.91 ± 3.65 | LYS311, NVAL306, LYS375 | LEU376, ILE308, PRO312 |
| | $COO^-$-q | 1 | −41.42 ± 3.73 | LYS311, NVAL306, LYS375 | PRO312, TYR310, ILE308 |
| | | 2 | −41.66 ± 3.39 | LYS311, NVAL306, LYS375 | ILE308, PRO312, LEU376 |

| | | | | | |
|---|---|---|---|---|---|
| | | 3 | −40.56 ± 3.18 | LYS311,<br>LYS375,<br>NVAL306 | PRO312,<br>ILE308,<br>TYR310 |
| | $O^-$-q | 1 | −43.25 ± 4.01 | LYS311,<br>NVAL306,<br>LYS375 | PRO312,<br>TYR310,<br>ILE308 |
| | | 2 | −40.15 ± 3.14 | LYS311,<br>NVAL306,<br>LYS375 | TYR310,<br>PRO312,<br>ILE308 |
| | | 3 | −40.82 ± 3.13 | LYS311,<br>NVAL306,<br>LYS370 | PRO312,<br>TYR310,<br>ILE308 |
| | Pristine | 1 | −21.58 ± 3.49 | LYS311,<br>CPHE378,<br>LYS375 | TYR310,<br>ILE308,<br>LYS311 |
| | | 2 | −22.42 ± 3.31 | LYS311,<br>CPHE378,<br>LYS375 | TYR310,<br>LYS311,<br>ILE308 |
| | | 3 | −18.74 ± 2.69 | LYS311,<br>CPHE378,<br>ASP314 | LYS311,<br>TYR310,<br>PRO312 |
| **Proto. int.** | $O^-$-half | 1 | −45.02 ± 3.60 | LYS340,<br>LYS331,<br>LYS321 | SER324,<br>HIS329,<br>GLY323 |
| | | 2 | −45.06 ± 3.56 | LYS340,<br>LYS331,<br>LYS321 | SER324,<br>ASN327,<br>HIS329 |
| | | 3 | −45.49 ± 3.40 | LYS340,<br>LYS331,<br>LYS321 | SER324,<br>HIS329,<br>GLY323 |
| | $S^-$-half | 1 | −65.36 ± 5.48 | LYS321,<br>LYS317,<br>LYS340 | GLY323,<br>SER324,<br>LEU325 |
| | | 2 | −69.75 ± 6.47 | LYS340,<br>LYS331,<br>LYS321 | SER324,<br>HIS329,<br>GLY326 |

| | | | | | |
|---|---|---|---|---|---|
| | | 3 | $-64.90 \pm 4.28$ | LYS340, | SER324, |
| | | | | LYS331, | ASN327, |
| | | | | LYS321 | HIS329 |
| | | 1 | $-66.93 \pm 2.65$ | LYS321, | GLY323, |
| | $COO^{-}$-half | | | LYS317, | CYS322, |
| | | | | LYS369 | GLY366 |
| | | 2 | $-78.16 \pm 7.27$ | LYS317, | THR319, |
| | | | | LYS321, | ASN368, |
| | | | | LYS370 | GLY367 |
| | | 3 | $-79.15 \pm 4.83$ | ARG349, | VAL350, |
| | | | | LYS353, | GLN351, |
| | | | | LYS347 | PHE346 |
| | | 1 | $-11.60 \pm 3.04$ | GLU338, | ASN327, |
| | Pristine | | | LYS340, | GLY326, |
| | | | | GLU342 | GLU338 |
| | | 2 | $-16.33 \pm 2.58$ | GLU342, | SER341, |
| | | | | LYS343, | GLU342, |
| | | | | LYS340 | LYS340 |
| | | 3 | $-25.12 \pm 3.72$ | GLU338, | ASN327, |
| | | | | LYS369, | GLY326, |
| | | | | LYS353 | HIS329 |
